# Engineering an in vitro retinothalamic nerve model

**DOI:** 10.1101/2024.03.06.582645

**Authors:** Giulia Amos, Stephan J Ihle, Blandine F Clément, Jens Duru, Sophie Girardin, Benedikt Maurer, Tuğçe Delipinar, János Vörös, Tobias Ruff

**Author notes:** Correspondence: Tobias Ruff, János Vörös.

## Abstract

Understanding the retinogeniculate pathway *in vitro* can offer insights into its development and potential for future therapeutic applications. This study presents a Polydimethylsiloxane-based two-chamber system with axon guidance channels, designed to replicate unidirectional retinogeniculate signal transmission *in vitro*. The system enables the formation of up to 20 identical functional retinothalamic networks on a single transparent microelectrode array. Using embryonic rat retinas, we developed a model where retinal spheroids innervate thalamic targets through up to 6 mm long microfluidic channels. We found that network integrity depends on channel length, with 0.5-2 mm channels maintaining over 90 % morphological and 40 % functional integrity. A reduced network integrity was recorded in longer channels. The results indicate a notable reduction in forward spike propagation in channels longer than 4 mm. Additionally, spike conduction fidelity decreased with increasing channel length. Yet, stimulation-induced thalamic target activity remained unaffected by channel length. Finally, we assessed the impact of stimulation frequency and channel length on the sustainability of the thalamic target spheroid response. The study found that a sustained thalamic calcium response could be elicited with stimulation frequencies up to 31 Hz, with higher frequencies leading to transient responses. In conclusion, this study shows how channel length affects retina to brain network formation and signal transmission *in vitro*.

## INTRODUCTION

Sensory systems form highly organised sensory maps in the brain during development. In the visual system, molecular guidance factors that are expressed along the optic pathway assure an initial retinotopic innervation of the thalamic target structures that are between 1-10 mm away (Schmitt et al. (2006); Triplett and Feldheim (2012); Leamey et al. (2007)). In later stages, spatiotemporally correlated activity waves in the retina refine those initially established connections (Faust et al. (2021); Ackman and Crair (2014)). Together those mechanisms preserve spatial sensory information and reduce redundant activity to maximize sensory resolution in the brain.

The developmental mechanisms that guide axons towards their target location and establish functional synaptic connections might be exploited to restore functional connections after damage. However, damaged retinal axons are unresponsive to signals present in the adult brain which translates to a limited regenerative capacity of retinal ganglion cells (Williams et al. (2020)). *In vivo* research suggests different approaches such as genetic modifications of the retinal ganglion cells (Yungher et al. (2015); Li et al. (2015); You et al. (2016)) or stimulated neuronal activity (Lim et al. (2016)) to enhance restoring retinal connections. Despite the potential of these approaches to understand the mechanisms of target innervation, the complexity and inaccessibility of the adult brain make purely animal-based *in vivo* research slow and expensive, often leading to inconclusive results with high variability.

*In vitro* neuronal systems have the advantage that multiple similar networks can be simultaneously exposed to the variable of interest resulting in higher efficiency and reproducibility. However, random *in vitro* cultures would not represent the directionality and typical axon length of the retinogeniculate pathway. To guide axons effectively *in vitro*, physical barrier-based axon guidance structures have mostly replaced purely surface patterned approaches (Yamamoto et al. (2023)) due to their long-term stability (Aebersold et al. (2016)). Polydimethylsiloxane (PDMS) based microfluidic structures are used to establish neuronal networks *in vitro* since they enable rapid prototyping of varying microstructure geometries and can be reliably mounted onto glass or microelectrode arrays (MEA) (Duru et al. (2022); Forró et al. (2018); Ihle et al. (2022); Mateus et al. (2022)). The guidance within those microstructures is based on the observation that axons are less likely to follow sharp turning angles and prefer growing along edges (Renault et al. (2016)). Exploiting this edge guidance principle, various network topologies have been developed in the past. Most designs implement 2 - 4 chamber systems that are separated by varying amounts of microchannels (Tong et al. (2021); Park et al. (2021); Moutaux et al. (2018); Isomura et al. (2015); Winter-Hjelm et al. (2023); Vakilna et al. (2021); van de Wijdeven et al. (2019); Pan et al. (2015); Chang et al. (2022); Brofiga et al. (2022)). To define pre- and postsynaptic neurons, some works implemented unidirectional channels between the chamber systems (Isomura et al. (2015); Winter-Hjelm et al. (2023); Vakilna et al. (2021)) or applied delayed seeding strategies (Tong et al. (2021); Moutaux et al. (2018)).

To develop new methods to functionally reinnervate the adult brain, we need *in vitro* model systems that allow us to screen for various parameters in a more efficient and reproducible way. These model systems should represent the characteristic directionality of the retinogeniculate pathway *in vivo*. Therefore, the goal of this work was to establish a basic *in vitro* functional retinogeniculate network model consisting of primary rat retinal and thalamic spheroids connected over unidirectional axon bundles of several mm length. To increase throughput of parameter screenings, the platform includes 10 - 20 identical networks on a single glass MEA. Based on the channel geometries of Forró et al. (Forró et al. (2018)), we have designed unidirectional two-node structures to promote the formation of unidirectional retinothalamic networks. We combined electrical stimulation and recording from glass MEAs with functional calcium imaging from primary rat thalamic spheroids to clearly separate pre-from postsynaptic activity. We investigated how the length of micrometer-sized PDMS-based axon guidance channels affects the directionality and fidelity of retinothalamic signal transmission. We demonstrate unidirectional spike transmission from the retina to the thalamus. Additionally, we reveal that while the length of the channel adversely impacts spike conduction fidelity, it does not influence stimulation-triggered thalamic activity. Lastly, we show that continuous stimulation of retinal axons at frequencies up to 31 Hz results in a prolonged calcium response in thalamic target spheroids.

## MATERIALS AND METHODS

### Design and fabrication of PDMS microstructures

All PDMS microstructures were designed using AutoCAD (Autodesk, San Rafael, USA) in a two-layer design. A template of each microstructure designed in the framework of this project can be found in the supplementary figures (Figures S2-S6). The first layer provides the axon guidance channels that are 4 µm high, while the second layer is 250 µm high and contains wells for cell seeding. Different channel lengths (0.5, 1, 2, 4, 6 and 8 mm) were chosen, while the other dimensions, such as channel cross-sectional area (8*×*4 µm), target and source node designs remained constant. The wafer and PDMS replica were fabricated using standard soft lithography by Wunderlichips GmbH (Zurich, Switzerland). The PDMS microstructures were designed to align with glass MEAs (60MEA500/30iR-Ti, Multi Channel Systems MCS GmbH, Reutlingen Germany) to stimulate, record and analyse signal propagation throughout the network.

### Preparation of microelectrode array

Glass MEAs with a 6*×*10 electrode grid, electrode spacing of 500 µm and an electrode diameter of 30 µm were used for experiments requiring stimulation. Standard 48-well plates (92048, TPP) were used for experiments without electrical stimulation.

The protocol for MEA preparation was adapted from Girardin et al. (Girardin et al. (2022)). On the date of substrate coating, the MEAs were rinsed with 1 % Sodium dodecyl sulfate (SDS) (L3771, Sigma-Aldrich), technical ethanol, technical isopropanol and ultrapure water (Millipore Milli-Q System, 18.2 MΩ *·* cm). The surfaces were blow-dried with a nitrogen gun and plasma cleaned for 2 min (18 W PDC-32G, Harrick Plasma, Ithaca, USA) at maximum power.

MEAs and 48-well plates were coated with 400 µl and 150 µl 0.1 mg/ml poly-D-lysine (PDL) (P6407, Sigma-Aldrich), respectively in phosphate buffered saline (PBS) (10010015, Gibco, Thermo Fisher Scientific) for 45 min at room temperature. The surfaces were rinsed three times with ultrapure water. 10 µg/ml laminin (11243217001, Sigma-Aldrich) in Neurobasal^TM^ Plus (A3582901, Gibco) was used as a secondary coating by adding the same volumes to the substrates and incubating at 37 °C overnight. Immediately before PDMS microstructure mounting, the laminin solution was rinsed with ultrapure water.

### PDMS microstructure mounting

The microstructures were soaked in 70 % ethanol for 15 min and dried for 15 min. The microstructures needed for the experiments were cut out of the PDMS replica using a surgical blade. All instruments used to handle the microstructures were submersed in ethanol for 15 min before usage and completely blow-dried with a nitrogen gun.

Following the microstructure preparation, a thin layer of ultrapure water was added to the coated MEAs so that the PDMS microstructures could glide freely on the glass surface. Using the forceps, one microstructure at a time was carefully transferred to a MEA and aligned to the electrodes under a benchtop microscope (SMZ-161, Motic). Following the alignment, residue water was carefully aspirated and the substrates were incubated at 37 °C for 45 min to enable maximum bonding of the microstructure to the glass surface. Once bonded, 1.5 ml of PBS was added to the substrates and the structures were placed in a desiccator to remove air trapped in the PDMS microchannels. The PBS was then replaced with retinal ganglion cell (RGC) medium (see Table 1 for medium composition) and the MEAs were covered with an autoclaved membrane cap. Small plastic dishes with PBS were added to the culture dish to reduce evaporation before placement in the incubator.

**Table 1.**
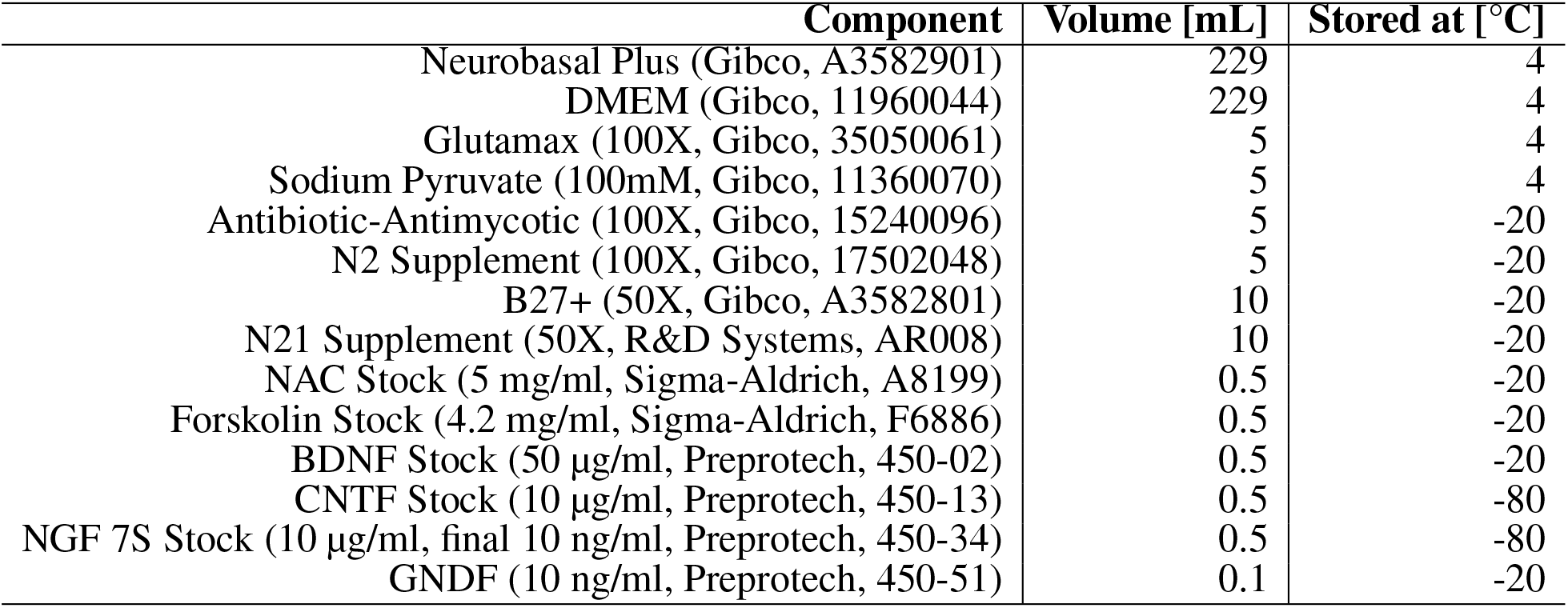
RGC medium composition. List of the components, volumes and storage temperatures for the RGC medium. The medium was used for the cocultures in all experiments.

| Component | Volume [mL] | Stored at [°C] |
| --- | --- | --- |
| Neurobasal Plus (Gibco, A3582901) | 229 | 4 |
| DMEM (Gibco, 11960044) | 229 | 4 |
| Glutamax (100X, Gibco, 35050061) | 5 | 4 |
| Sodium Pyruvate (100mM, Gibco, 11360070) | 5 | 4 |
| Antibiotic-Antimycotic (100X, Gibco, 15240096) | 5 | -20 |
| N2 Supplement (100X, Gibco, 17502048) | 5 | -20 |
| B27+ (50X, Gibco, A3582801) | 10 | -20 |
| N21 Supplement (50X, R&D Systems, AR008) | 10 | -20 |
| NAC Stock (5 mg/ml, Sigma-Aldrich, A8199) | 0.5 | -20 |
| Forskolin Stock (4.2 mg/ml, Sigma-Aldrich, F6886) | 0.5 | -20 |
| BDNF Stock (50 µg/ml, Preprotech, 450-02) | 0.5 | -20 |
| CNTF Stock (10 µg/ml, Preprotech, 450-13) | 0.5 | -80 |
| NGF 7S Stock (10 µg/ml, final 10 ng/ml, Preprotech, 450-34) | 0.5 | -80 |
| GNDF (10 ng/ml, Preprotech, 450-51) | 0.1 | -20 |

### Cell culture

Primary thalamic and retinal cells from E18 embryos of pregnant Sprague-Dawley rats (Dissected at EPIC, ETH Phenomics Center, Switzerland and imported from Janvier Labs) were used for all experiments. Compliance with 3R regulations was ensured (Russell and Burch (1959)). Previous to the experiments, approval was obtained from the Cantonal Veterinary Office Zurich, Switzerland, under license SR 31175 - ZH048/19. Briefly, E18 time-mated pregnant rats were sacrificed and the embryos were removed. For retina dissection from the embryonic eye, the lens and hyaloid vasculature were gently separated from the retina using a sharp pair of forceps under the benchtop microscope. Embryonic retina and thalamus tissue were dissected and stored in Hibernate medium on ice until dissociation. Cells were dissociated as described previously (Ihle et al. (2022)).

### Spheroid creation and seeding

Commercially available microwell plates (AggreWell^TM^ 400 24-well plate, 34415, StemCell Technologies) were used to generate retinal and thalamic spheroids (250, 500 and 1000 cells/spheroid). To prevent adherence, the microwells were coated with 500 µl of AggreWell^TM^ Anti-adherence rinsing solution (7010, StemCell Technologies) and rinsed with 2 ml Neurobasal^TM^ Plus (A3582901, Gibco) preheated to 37 *^◦^*C. The medium was replaced with 1 ml RGC medium per well and the microwell plate was left in the incubator at 37 °C for the medium to reach an appropriate pH and temperature value.

The spheroids were fluorescently labelled depending on their use in the experiment through adeno-associated virus (AAV) mediated transduction. Retinal source spheroids were transfected with mRuby virus (mRuby, scAAV-DJ/2-hSyn1-chl-mRuby3-SV40p(A)), while thalamic target spheroids were labelled using GCaMP8m virus (ssAAV-DJ/2-hSyn1-jGCaMP8m-WPRE-SV40p(A)) or EGFP virus (GFP, V-DJ/2-hSyn1-chl-EGFP-SV40p(A)). All adeno-associated viral vectors were provided by the Viral Vector Facility of the University of Zurich, Switzerland. The GCaMP8m transduction was repeated after five days *in vitro* (DIV) to achieve stronger fluorescence. Cell solution volumes for 250, 500 and 1000 cell spheroids were added to the microwells. Lastly, the microwell plate was centrifuged at 100 g for 3 min and transferred to 37 *^◦^*C.

After one day in the microwells, the retinal spheroids were manually seeded into the source inlets of the PDMS microstructures using a 10 µl pipette. Thalamic spheroids were added to the target inlets after five or twelve days to reduce the number of back-growing target axons. For Sholl analysis, single retinal spheroids of 250, 500 and 1000 cells were additionally transferred to a central position in each well of a 48-well plate using a 10 µl pipette. Half of the culture medium was exchanged every three to four days. In all experiments, the day of source neuron seeding was considered as DIV zero.

### Imaging

The neuronal cultures were imaged with a confocal laser scanning microscope (CLSM, FluoView 3000, Olympus) maintaining cell culture conditions (Pecon). Depending on the experiment, one to three channels were acquired in combination with phase contrast brightfield images: 488 nm (GCaMP8m or Alexa Fluor 488 antibody), 561 nm (mRuby), 594 nm (Alexa Fluor 594 antibody) and 640 nm (Alexa Fluor 633 antibody). A 10x (UPFLFLN10XPH, NA=0.3) was used for tilescan imaging and a 20x (UPLFLN20XPH, NA=0.5) was used for Calcium imaging.

For visual representation, CLSM images were processed using the open-source software Fiji (Schindelin et al. (2012)). Logarithmic processing was applied to the pixels of the images in Figures 1C, 1D, 2Aii, S1A and S1B to increase visibility for presentation purpose.

**Figure 1.**
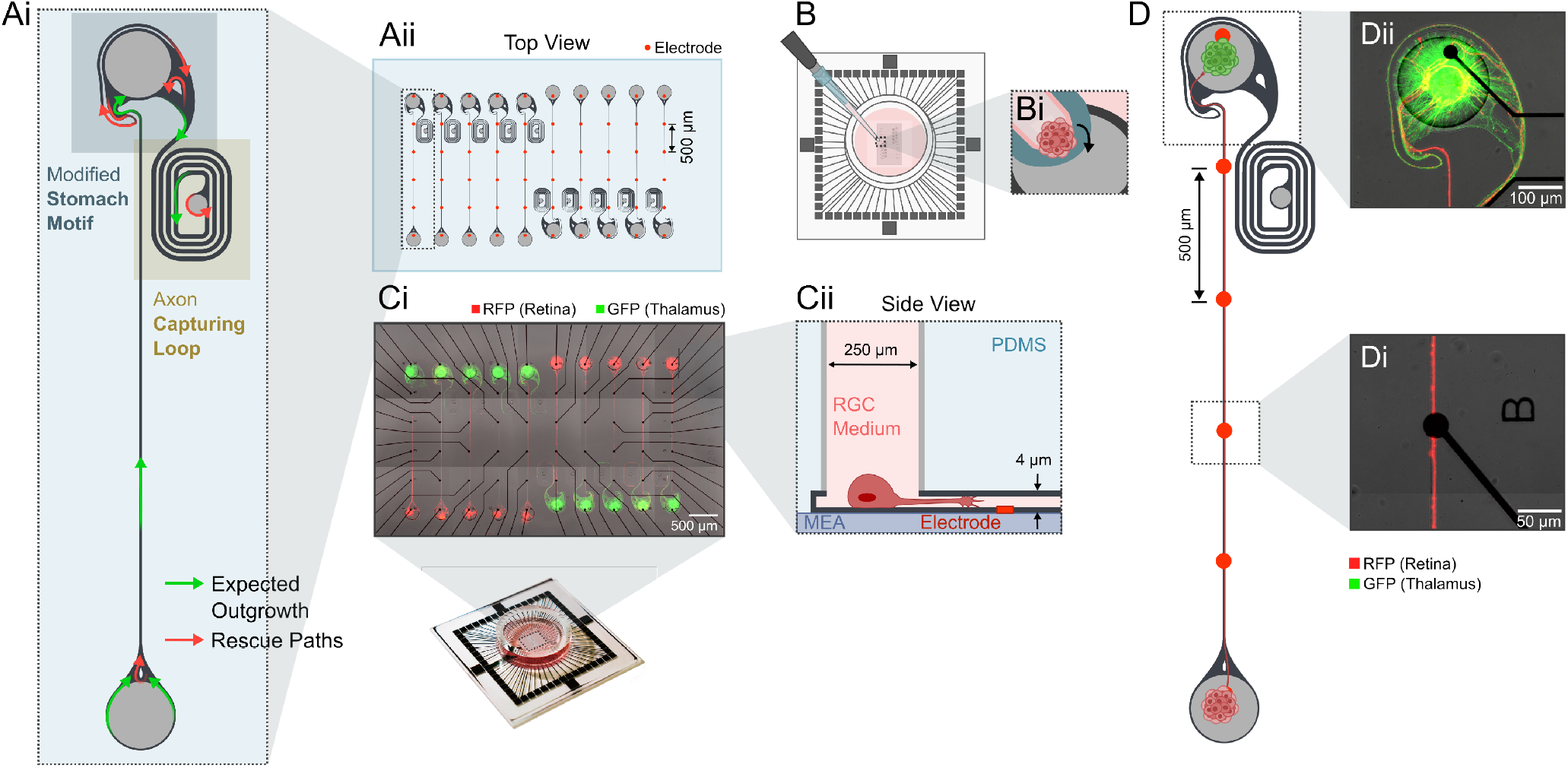
Experimental design to create unidirectional retinothalamic networks on a glass MEA. **(Ai)** The architecture of the microstructure favours directionality by a modified stomach structure motif (Forró et al. (2018)) and an axon capturing loop that traps thalamic axons into a loop preventing them from growing back into the retinal source chamber. **(Aii)** Example CAD layout of the 250 µm thick PDMS microstructures with 2 mm long axon guidance channels that align with a 60 electrode MEA. **(B)** PDMS microstructures are aligned on top of the glass MEA using a thin layer of water. **(Bi)** After drying, RGC medium is desiccated into the microstructure and spheroids are individually seeded into the different seeding wells. **(Ci)** Retinal spheroids (rat, E18.5,mRuby, red), that are seeded into the source well, grow axons towards a target seeding well containing thalamic spheroids (rat, E18.5, EGFP, green). As *in vivo* only retinal ganglion cells (RGCs) form long axonal projections to the brain we assume that axons within the microchannel are from RGCs. **(Cii)** The wells contain the cell bodies in their target location. **(D)** Example of one retinothalamic network where **(Di)** axons align with the electrodes and **(Dii)** RGC axons innervate the thalamic target spheroid.

### Immunofluorescence staining

To characterise the axonal outgrowth in 250, 500 and 1000 cell retinal spheroids, the samples in the 48-well plate were immunostained and imaged with a CLSM at DIV 5. The spheroids were fixed with 4 % paraformaldehyde (PFA) (1004960700, Sigma-Aldrich) for 20 min at room temperature. The PFA was carefully removed under the chemical hood, followed by two washing steps with PBS. The cells were then permeabilised using PBS containing 0.1 % Triton X-100 (9002931, Sigma-Aldrich) and 1 % BSA for 1-1.5 h. The medium was exchanged with a primary PBS antibody solution containing 0.1 % Triton-X, 1 % BSA and the primary antibodies: mouse anti-neurofilament (SMI 312, 837904, Biolegend) and rabbit anti-MAP2 (PA5-17646, Invitrogen, Thermo Fisher Scientific) in a 1:2000 solution. After approximately 1.5 h, the samples were rinsed twice with PBS and 1 % BSA before incubating them with the secondary antibody solution overnight at 4 °C. The secondary antibody solution was composed of 1 % BSA, Alexa Fluor 488 goat anti-rabbit antibody (1:2000, A32731, Invitrogen, Thermo Fisher Scientific) and Alexa Fluor 633 goat anti-mouse antibody (1:2000, A21050, Life Technologies) in PBS. The next day, samples were washed twice with PBS and left in PBS until imaging.

Images were analysed using a Fiji plug-in called NeuriteJ (Torres-Espín et al. (2014)). NeuriteJ was initially developed to investigate neurite arbors in organotypic cultures and provides a semi-automated method to quantify axonal growth from any defined central body (Sholl (1953)). Axonal growth is quantified by creating a series of concentric shells of increasing radii around the central body and counting the number of intersections of the arbor with these shells (Torres-Espín et al. (2014)).

Concentric circles were drawn from the edges of the spheroid bodies at 25 μm intervals. The number of intersections at each distance was plotted for every spheroid and the maximum number of intersections was determined (Figure S1). Only distances greater than 300 μm were considered in the analysis to reduce false-positive intersection counts caused by the non-specific binding of antibodies to dendritic outgrowths. The results were normalised by dividing the maximum number of intersections by the mean of the 250 cell spheroid count of the respective experimental repetition. Spheroids were excluded from the analysis if they adhered to the wall of the well, preventing the formation of a circular neurite arbor, or if more than one spheroid was seeded into the same well.

### Electrophysiology

A MEA2100-Mini headstage (MEA2100-Systems, Multi Channel Systems MCS GmbH, Reutlingen, Germany) was used to record electrical activity from the two-node neuronal networks at DIV 7, DIV 14, DIV 21 and DIV 28. The MEA headstage was placed inside the preheated (37 °C) CLSM incubator at 5 % V/V CO_2_ in air. The humidity was not controlled.

The cultured neuronal networks were left in the CLSM incubator for at least 3 min to settle before starting the recording. After a 3 min adaption phase within the CLSM, data was acquired for 1 min from all 60 electrodes simultaneously at a sampling frequency of 25 kHz. Additionally, a CLSM image was taken to assess neurite outgrowth.

Networks were only included in the analysis if a visual inspection of the CLSM images at DIV 28 showed that (1) the spheroids stayed in their respective well, (2) the axon bundle did not escape from the microchannel, (3) the full width of the microchannel was over the electrodes, (4) the network had no neuritic growth that connected it to a different network on a MEA and (5) the axon bundle reached the first electrode at any stage during the four weeks. A network was excluded from any subsequent analysis if one or multiple of these criteria were violated.

Raw data of the spontaneous electrical activity were filtered with a high-pass second-order Butterworth filter with a cut-off frequency of 200 Hz. For each electrode, the baseline noise of the signal was calculated using the median absolute deviation (MAD). The MAD is less susceptible to outliers in the voltage trace and, consequently, action potentials in the spike traces have less influence on the noise estimation (Ihle et al. (2022)). Hence, using the MAD to calculate the standard deviation is a more robust approach than calculating the standard deviation directly from the spike trace. The standard deviation σ from the MAD was defined as follows (Leys et al. (2013)):

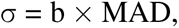

where b = 1.4826 is a constant and the MAD is calculated as

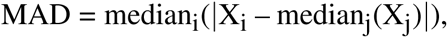

with X_i_ being the i^th^ data point of the raw voltage trace. Spikes were detected by identifying negative voltage peaks below a threshold of five times the calculated standard deviation σ. Successive peaks were discarded to avoid duplicates if they occurred within 3 ms from a detected spiking event.

The network formation and signal transmission in the microstructures were evaluated based on eight metrics derived from the spike trains and CLSM images, namely the percentage of intersected electrodes per network, the morphological network integrity, the percentage of active electrodes per network, the functional network integrity, the mean firing rate, the conduction speed, the conduction fidelity and the percentage of forward propagating spikes. The computation of each metric is explained in supplementary Table S1. Only morphologically intact and active networks were included into subsequent analyses if not stated otherwise.

### Calcium imaging

To further validate and compare the directional and functional connectivity in the two-node neuronal networks, the activity in the target was measured using GCaMP8m while electrically stimulating the axon bundle in a microchannel. The most proximal electrode was used for stimulation. The electrical stimuli used in the experiment had square wave shapes (first cathodic, then anodic segment) with a fixed peak-to-peak voltage of 2 V and a pulse width of 200 µs.

At DIV 21, the response of the cultures to continuous stimulation was assessed. The stimulation protocol consisted of two sections of 15 s each. In the first 15 s, the spontaneous fluorescence dynamics were measured to quantify the baseline activity. This was followed by 15 s of continuous stimulation by a pulse train. The applied inter-stimulus interval (ISI) of the pulse train was variable and ranged from 4 ms to 256 ms. Each ISI variation was applied six times in succession to each network, whereby the order of ISI variation was randomised.

At DIV 28, the response to a stimulus train of 16 stimuli with a fixed ISI of 5 ms was assessed. The spontaneous fluorescence dynamics in the target were measured for 200 ms before applying the stimulus train, followed by a 3 s break to allow the network to recover to baseline levels. The cycle was repeated 40 times for each neuronal network.

During recording, MEAs were kept at 37 °C and 5 % CO_2_. Stimulation was started after a minimum of 3 min to give the culture time to settle. The fluorescence dynamics of GCaMP8m in the thalamic spheroid during continuous stimulation at DIV 21 were recorded at 10 ms per frame, whereby ten subsequent frames were averaged. The fluorescence dynamics at DIV 28 were recorded with a frame rate of 10 ms per frame without averaging. For both stimulation protocols, the 488 nm laser at 1 % was used with a 20 *×* objective (UPLFLN20XPH, NA=0.5).

In post-processing, region of interests (ROIs) were drawn with Fiji around each spheroid to exclude background regions with no informational content. The mean grey value F0 of each ROI was calculated for the baseline fluorescence levels. The normalised fluorescence change dF/F0 was defined as

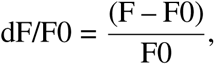

for each frame. The data was further processed as described below for the two stimulation protocols.

### Processing of continuous stimulation recordings

The slope of the normalised fluorescence trace during continuous stimulation was calculated by fitting a linear function to the region between the peak evoked response (PER) dF/F0_max_ in the first 5 s after stimulation onset and the end of the stimulation cycle. The mean slope m of the linear function was calculated over six cycles.

### Processing of stimulus response recordings

The PER was defined as the maximum fluorescence change after stimulation dF/F0_max_. The mean and variance of the PER were calculated over 40 cycles.

Additionally, the transmission fidelity, defined as the percentage of detectable stimulus responses after axon bundle stimulation, was computed. A stimulus response was classified as detectable if the PER exceeded the baseline fluorescence activity by a factor of five standard deviations. The standard deviation was calculated using the MAD.

### Statistical Analysis

Statistical analyses were performed with the GraphPad Prism software (version 9.4.0, GraphPad Software, San Diego, USA). The data was tested for normal (Gaussian) distribution using normal quantile-quantile plots and the D’Agostino & Pearson test (D’Agostino (2017)). If the normality assumption could not be rejected (p-value of D’Agostino & Pearson test *>* 0.05 and quantile-quantile plot displaying a roughly straight line), the population was considered consistent with a Gaussian distribution.

If the data did not deviate from normality, a one-way ANOVA (Fisher (1992)) was used to check for significant effects in the data. If ANOVA showed a significant effect, a post-hoc test was performed using Tukey’s correction for the multiple comparison problem (Keselman and Rogan (1977)).

If the population deviated from normality, the distribution-independent Kruskal-Wallis test (Kruskal and Wallis (1952)) was used for statistical evaluation of non-paired and Friedman’s test (Friedman (1937)) for paired data sets. Both tests are non-parametric and test the null hypothesis that the compared groups have the same underlying distribution. If the null hypothesis was rejected in the Kruskal-Wallis or Friedman’s test (p *≤* 0.05), Dunn’s post-hoc test (Dinno (2015)) was used to identify which pairwise groups in the dataset show significant differences in their distribution. Calculating the p-value with Dunn’s multiple-comparison procedure corrects for the number of comparisons made.

The comparisons made are listed in the result section. Differences were considered significant as described in the text.

## RESULTS

### Directional axon guidance elements promote functional retinothalamic network formation with up to 6 mm long axons

To model functional long-distance retinogeniculate signal transmission *in vitro*, we designed a PDMS-based two-chamber system that is connected by an axon guidance channel (Figure 1Ai, Aii). We implemented a directional target well that promotes unidirectional axon growth (Figure 1Ai) in order to achieve unidirectionality as seen in the retinogeniculate circuits *in vivo*. The target well includes axon guidance elements from previously published work (Forró et al. (2018)) and an axon guidance spiral to capture retinal and thalamic axons with the intend to reduce the probability of axons growing backwards. Both seeding wells are 250 μm wide and deep and enabled the seeding of spheroids with up to 1000 neurons (Figure 1B, Bi, Ci, Cii). The 4 *×* 10 μm small cross section of the axon guidance channels prevents migration of neuronal somas and thereby assure long-term morphological integrity of our system (Figure 1Ai). 10-20 individual retinothalamic networks can be established on a single 60 electrode glass MEA in which the axon guidance channels align with the stimulation and recording electrodes (Figure 1Ci). For the retinothalamic nerve model we used primary dissociated embryonic rat retinas. To identify an appropriate spheroid size that will generate a sufficient number of axons for the *in vitro* model, we assessed the axonal outgrowth of spheroids ranging from 250 - 1000 cells/spheroid (Figure S1). Based on results from Sholl analysis (Sholl (1953)) we decided to use retinal spheroids of 500 cells/spheroid for all experiments presented in this work. Our results show that retinal spheroids adhere to the surface (Figure 1D) and extend axons that are guided across the stimulation electrodes (Figure 1Di) to innervate the thalamic target spheroid (Figure 1Dii) within just 7 days.

In adult rats, retinal axons need to extend about 2 cm to reach the primary visual centers such as the superior colliculus in the brain (Abbott et al. (2013)). Rebuilding a retinothalamic nerve model with long axon guidance channels enables us to understand how retinal signal transmission is affected by distance. Thus, we varied the channel length between 0.5-8 mm (Figure 2Ai) and asked how the spatial constrain within 4 *×* 10 µm small channels of varying channel length affects the morphological (Figure 2Aii) and functional integrity of the mRuby labelled retinal axons over time (1-4 weeks *in vitro*). We assessed the growth of axons in the microchannels by quantifying the percentage of electrodes that were intersected with retinal axons (mRuby fluorescence). Our results show that the average percentage of axon-intersected electrodes decreases from about 80-100 % for the 0.5-2 mm long channels to below 50 % for the 8 mm channels. The culture age, however, did not significantly influence the percentage of intersected electrodes indicating that within DIV 7, axons reached their maximum extension (Figure 2 B). The percentage of active electrodes (spike rate above 1 Hz) was lower at around 50 % for the shorter channels (0.5-2 mm) and dropped to below 20 % for the 8 mm channels (Figure 2Bi). Interestingly, the percentage of active electrodes dropped within 3 weeks to below 10 % for the longer 6-8 mm channels (Figure 2Bi).

**Figure 2.**
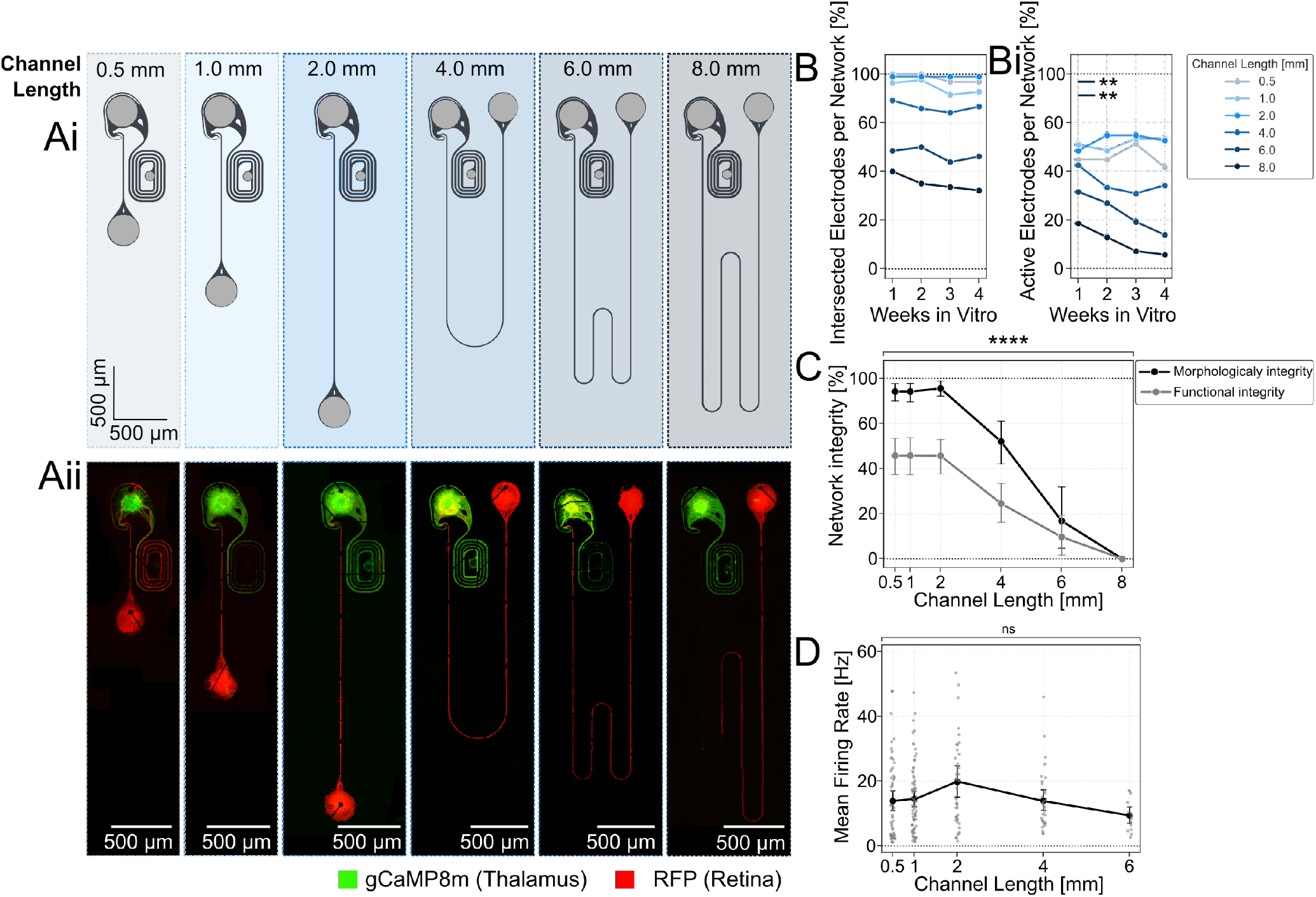
Retinal spheroids form morphological and functional intact networks with axons that have connection paths of up to 6 mm on 60 channel glass MEAs. **(Ai)** Microstructure design of the two-node structures with an increasing length of axon guidance channels. Light grey areas represent seeding wells that are open from the top. Dark grey regions are 4 μm tall microchannels. **(Aii)** Fluorescently labelled retinal neurons (AAV2/DJ-mRuby) reach the thalamic target spheroids (GCaMP8m) through up to 6 mm long channels at DIV 21. **(B)** Fluorescent imaging and electrical recording of spontaneous neuronal activity along the axon guidance channel were used to assess the neuronal networks. The average number of electrodes on the MEA that are intersected with fluorescently labelled axons and the **(Bi)** average number of active electrodes per network was measured over time and compared for different channel lengths. A decline in active electrodes per network was observed in networks with 4, 6 and 8 mm long axon channels. **(C)** The morphological and functional network integrity declines significantly for channels longer than 2 mm. A network was considered morphologically intact when red fluorescing retinal axons (AAV2/DJ-mRuby) were detected up to the target well. Networks were considered functionally intact when spontaneous neuronal activity above 1 Hz was measured along the entire channel. ****: p *≤* 0.0001 (Kruskal-Wallis test). **(D)** A similar spontaneous activity rate per active electrode could be measured for channels up to 6 mm in length. Inactive electrodes (spike rate below 1 Hz) were excluded from the measurement.

The platform design enabled to culture 10-20 networks in parallel to increase throughput of parameter screening. Results showed that platforms with 0.5-2 mm long channels had more than 40 % functionally and more than 90 % morphologically intact networks (Figure 2C). For channels exceeding 2 mm there was a significant decline in both morphological and functional network integrity (Figure 2C) (Kruskal-Wallis test (****: p *≤* 0.0001)). At a channel length of 8 mm, no networks remained fully intact, neither morphologically nor functionally (Figure 2C). At a 6 mm length, approximately 10 % of the networks on a platform managed to maintain their integrity. Despite the observed decline in intact networks, the mean firing rates revealed no significant differences across channels up to 6 mm in length (Figure 2D). Only functionally intact networks where included. This finding proves that the extension of axonal growth challenges network integrity.

### The retinothalamic nerve model promotes unidirectional spike propagation

*In vivo* spiking information is unidirectionally transferred from the retina into the brain. To model the unidirectional information transfer *in vitro*, we (1) implemented unidirectional guidance elements into the thalamic target well and (2) seeded the thalamic target spheroids at DIV 5. This gives retinal axons an advantage to grow first and reduces the axonal growth potential in the thalamic target spheroids. We assessed the directionality of spontaneously evoked spikes along the axon guidance channel within the retinogeniculate neuronal networks. (Figure 3Ai and ii). The directionality of the spikes was assessed by correlating spike latencies (Figure 3B) after the detection of a spike in the green electrode (closest to target well) (Figure 3Ai) or blue electrode (closest to source well) (Figure 3Ai). Results show that more than 85.9 % of the spikes are travelling from the retinal spheroids towards the thalamic targets (forward propagation). However, the average percentage of forward propagating spikes per network shows a significant decrease in 4 mm and 6 mm long channels (Figure 3C). We have also measured the spike conduction velocities over time in channels of different lengths to assess the functional maturation of the retinal axons. The average spike conduction speed over all measured DIVs significantly declined from 0.81 m/s for the 1 mm long channels to 0.67 m/s for the 6 mm long channels (Figure 3D). In line with previous publications the average spike conduction speed showed a significant increase from 0.6 m/s to 1 m/s between DIV 1 and 4 weeks across all channel lengths (Figure 3E). Interestingly, the increase of conduction speed was strongest in the shorter channels (1 mm and 2 mm) and decreased in the 4 to 6 mm long channels (Figure 3F).

**Figure 3.**
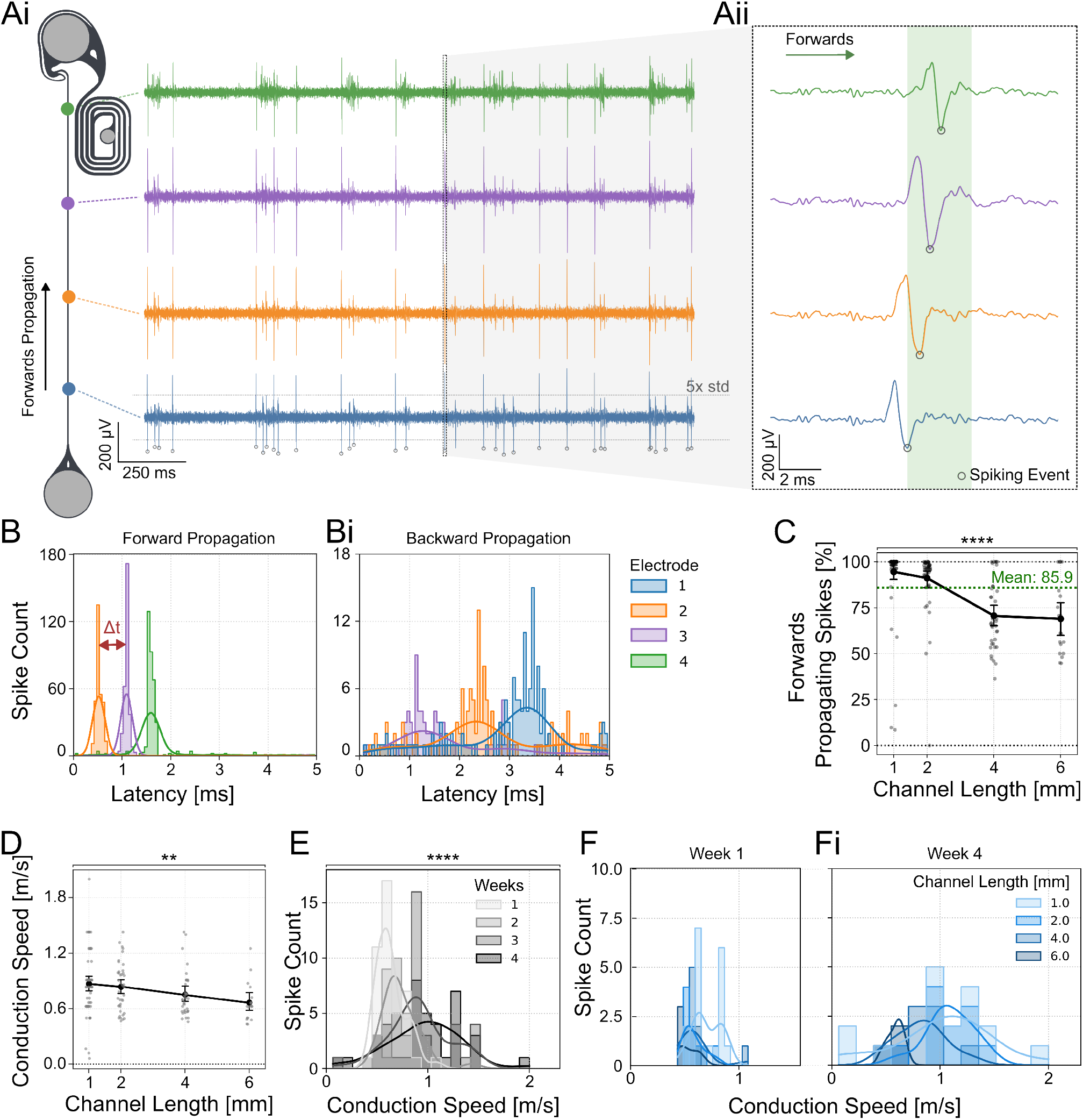
Directionality of axonal spike propagation within PDMS microchannels. **(Ai)** The directionality and speed of neuronal spikes were determined by the spike time differences measured at the electrodes aligned with the PDMS channel. **(Aii)** Voltage traces of spikes at different electrodes along the PDMS microchannel. Example of a forward propagating spike starting at the blue electrode. The spike time was determined using the negative peak of the spike waveform. **(B)** STTH of forward and **(Bi)** backward propagating spikes within a 5 ms time window which was used to assess directionality in C. The plot illustrating forwards propagation was triggered on the most proximal electrode (blue), while the plot showing backwards propagation was triggered on the most distal electrode (green) to show the spike propagation through the PDMS channel. **(C)** The directionality of spikes appearing closest to the source and target location was determined by measuring correlated activity in the other electrodes of the same channel. Each data point represents one retinothalamic network. The percentage of forward propagating spikes shows a significant decline with increasing channel lengths (Kruskal-Wallis test: p*<*0.0001). The percentage of forward propagating spikes is significantly different between 1 and 2, 1 and 3, 1 and 6 and 2 and 4 mm long channels (Dunn’s multiple comparison test) (Table S8.) **(D)** The average spike conduction speed was measured along the channels at DIV 28. Each data point represents the average spike propagation speed of all spikes measured within a 1 minute time window of one retinothalamic network. A decline in conduction speed between 1 and 6 mm long PDMS channels was observed (Kruskal-Wallis, test:p=0.0059, Dunn’s post-hoc analysis: p=0.0121). **(E)** Histogram of the spike velocities over all channel lengths show a significant increase in conduction speed within the first 4 weeks in cultures (Kruskal-Wallis test: p*<*0.0001). **(F)** Comparison of the spike conduction speed of axons in different channel lengths. At week 1 the channel length has no effect. **(Fi)** At week 4 the shorter channel lengths (1 mm and 2 mm) show the highest increase in spike speed. The spike count was recorded within 1 minute

### Channel length negatively affects spontaneous spike conduction fidelity but not stimulation induced target activity

In rats retinal axons need to reach their neuronal target up to 2 cm away and maintain a reliable signal transmission. Here, we emulated the formation of nerve like structures by physically confining axons within a small PDMS microchannel of a 4*×*10 µm cross-section. We asked how the physical confinement of channels having a length of up to 8 mm will affect the spontaneous spike propagation and stimulation fidelity in the thalamic target neurons. We assessed the spike conduction fidelity of forward (retina to thalamus) moving spikes by quantifying how many spikes measured at the first electrode (closest to the retinal spheroids) reached the electrode closest to the thalamic target spheroid (Figure 4A). Our results show that the spike conduction fidelity of forward propagating spikes significantly dropped with increasing channel lengths from close to 50 % for 1 mm long channels down to approximately 25 % for 6 mm long channels (Figure 4B). To measure changes in GCaMP fluorescence, we integrated the microelectrode array stimulation unit into a CLSM equipped within a CO_2_ 37*^◦^*C incubation chamber (Figure 4C). To assess how stimulation of the retinal axons affects thalamic target activity we transfected the thalamic neurons with the functional calcium indicator GCaMP8m (Figure 4D). Our results show that stimulation of retinal axons was able to modulate thalamic target activity (Figure 4F, G); and that the transmission fidelity of target stimulation in morphologically intact networks was independent of the channel length (Figure 4E).

**Figure 4.**
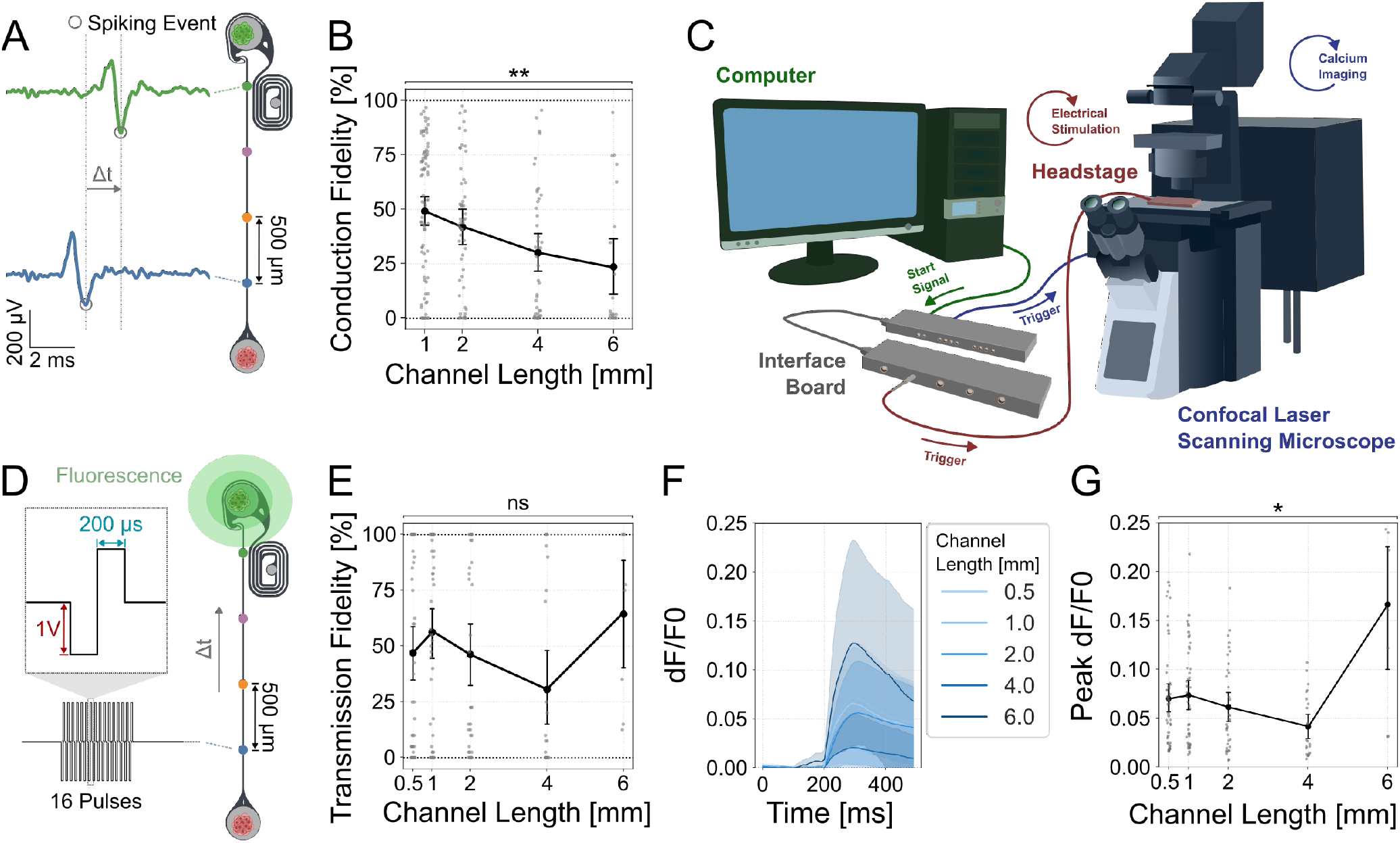
Spike conduction fidelity decreases with increasing channel lengths but does not significantly affect stimulation induced transmission fidelity of the thalamic target tissue **(A)** The percentage of spontaneous forward propagating spikes that can be measured at both (blue and green electrodes) vs. only at the source (blue electrode) were measured to quantify conduction fidelity. **(B)** Quantification of the spontaneous spike conduction shows a significant decline in the conduction fidelity (Kruskal Wallis test, p*<*0.0019). Dunn’s multiple comparison test identified significant differences between 1 and 4 mm (adjusted p*<*0.0182) and 1 and 6 mm (adjusted p*<*0.0107) channel lengths. Error bars are SEM. **(C)** Setup combining electrophysiological recording and stimulation with functional calcium imaging using a confocal laser scanning microscope. **(D)** To measure the transmission fidelity, retinal axons were stimulated with 40 cycles of 16 pulse bursts at 250 Hz at the source (blue electrode) and the corresponding thalamic response was measured using functional calcium imaging (GCaMP8m, ssAAV-DJ/2-hSyn1-jGCaMP8m-WPRE-SV40p(A)). **(E)** Quantification of the transmission fidelity shows no significant difference between the different channel lengths. **(F)** Stimulation induced calcium traces of thalamic spheroids. Shaded areas are 95 % confidence intervals. In all graphs, each data point represents one retinothalamic network. **(G)** Quantification of the peak evoked calcium response for different channel lengths (KW: p=0.0143).

### Axonal stimulation frequencies of up to 31 Hz at the retinal source elicits a sustainable thalamic response

Next, we asked how the stimulation frequency and channel length affects the sustainability of the thalamic target spheroid response. We, therefore, stimulated the retinal axons at DIV 21 for 15 s with increasing stimulation frequencies and measured the corresponding GCaMP response as dF/F0 of the thalamic target spheroids (Figure 5A, B, S7). Thalamic responses could be elicited in networks with up to 4 mm long channels (Figure 5A). We quantified the response sustainability by measuring the slope of the calcium response decline from its peak value until the end of the stimulation cycle (Figure 5C). Our results show that the thalamic calcium response remains at a steady level for stimulation frequencies up to 32 Hz (Figure 5D). For higher stimulation frequencies (62 - 250 Hz) we observed a transient increase in GCaMP fluorescence that initially surpassed the sustained response but decreased back to sustained levels of the 32 Hz stimulation within about 7 s (Figure 5C, D). Additionally, the results indicate that the maximum stimulation frequency that induces a sustained response is independent of the channel length (Figure 5A, D). For the highest stimulation frequencies (125 and 250Hz), however, a shorter channel resulted in a higher transient peak evoked response. Consequently lower channel lengths had a steeper decline of the transient response back to base levels (Figure 5D, S7B).

**Figure 5.**
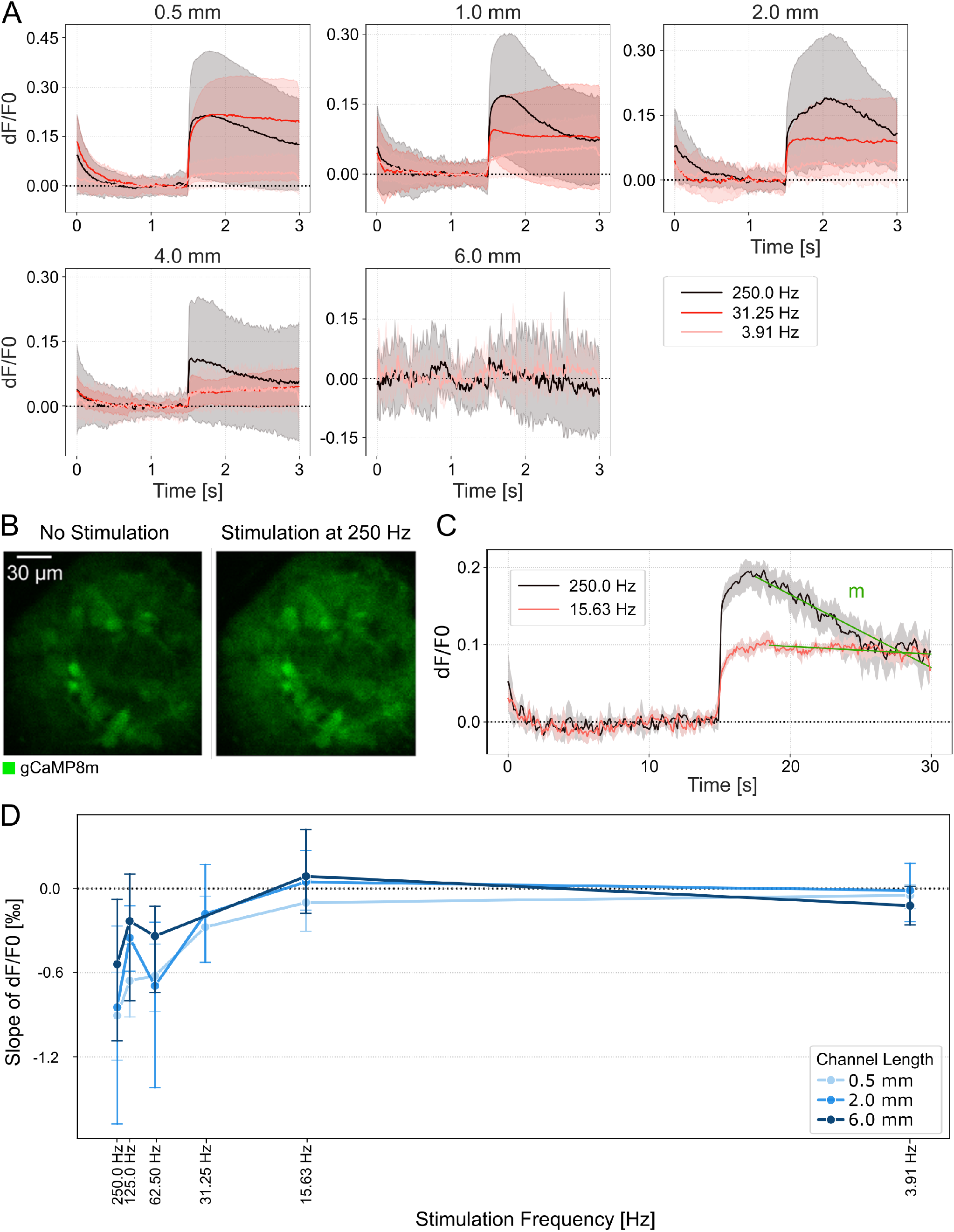
Continuous stimulation of thalamic target sheroids elicits a sustained calcium response. **(A)** Average calcium traces of thalamic spheroids in response to axonal stimulation at different frequencies over increasing channel lengths. Stimulus onset at 15 s. The areas represent the estimated mean with a <u>95 % confidence interval.</u> **(B)** Fluorescence images of a thalamic spheroid before and during electrical stimulation at 250 Hz. **(D) (C)** Example calcium traces of a sustained (red) and transient (black) response recorded at 10ms/frame. **(D)** Average calcium response decay during continuous stimulation. Negative values indicate a negative slope and thus a decrease in fluorescence intensity over the stimulation period. A value close to 0 indicates a sustained response. Error bars, 95 % confidence interval.

## DISCUSSION

In this study, we aimed to replicate the retinothalamic signal transmission pathway *in vitro* on a microelectrode array for electrical stimulation and recording. We emulated the pathway using a unidirectional PDMS-based two-chamber microstructure that is connected with 10*×*4 μm narrow axon guidance channels of varying lengths. Using primary rat retinal and thalamic spheroids, we asked how the channel length affects network integrity and signal transmission fidelity. Our results showed that (1) functional networks can be created with up to 6 mm long channels, (2) the directional target well in combination with a delayed target seeding strategy resulted in directional networks and (3) that stimulation of retinal axons within the PDMS channel could reliably activate thalamic target spheroids.

### Modeling the retinothalamic pathway

To our knowledge this is the first model combining retinal and thalamic spheroids into a network on a microelectrode array. This is especially important when modeling the network dynamics of an *in vivo* neuronal network *in vitro*. Previous works using cortical and hippocampal neurons have shown that neuronal identity influences network behavior (Vakilna et al. (2021); Chang et al. (2022); Brofiga et al. (2022); Virlogeux et al. (2018)) suggesting that different neuronal types should be included to accurately model *in vivo* networks *in vitro*. In this work, we used dissociated retinas from E18 rat embryos. At that time, retinas have developed most of their RGCs and contain horizontal and amacrine cells (Rapaport et al. (2004); Whitney et al. (2023)). At this stage, only RGCs develop axons. Therefore, we did not purify retinal cells prior to seeding. In E14 embryonic mouse retinas, RGCs represent about 14 % of the total cell population. Assuming a similar percentage in rat embryonic retinas results in an optimistic estimate of about 70 RGCs per spheroid within our culture system. This number matches the average number of maximum intersections measured in the Sholl analysis (Figure S1) (Mcloughlin et al. (2023)).

### Random vs. defined neuronal networks

Up to 20 identical retinothalamic networks could be aligned onto a single MEA, which is higher than previously published *in vitro* neuronal networks using different cell types (Moutaux et al. (2018); Virlogeux et al. (2018); Winter-Hjelm et al. (2023); Vakilna et al. (2021)). Moreover, our strategy resulted in a comparatively high number of functional networks (Ihle et al. (2022); Duru et al. (2022)). The vast majority of studies investigating neuronal networks in PDMS microstructures used suspended cells for seeding (Ihle et al. (2022); Girardin et al. (2022); Jungblut et al. (2009); Holloway et al. (2019); Hong et al. (2016)). While these results are not directly comparable due to the different cell types used, the high percentage of active networks in our structures might be attributed to the use of spheroids instead of cell suspensions. Moreover, the use of spheroids allowed us to selectively seed different cell types in target and source wells and reduced the amount of cells required for our experiments.

### Spontaneous activity

In primary neuronal cultures the spontaneous firing rate increases as the networks mature. In primary rat hippocampal neuronal networks spontaneous activity increases between DIV 4 and DIV 10-14 before reaching a plateau (Winter-Hjelm et al. (2023)), probably indicating maturation and synapse formation to be complete after DIV14 (Chiappalone et al. (2006); Ichikawa et al. (1993). The spontaneous activity within the here described retinothalamic networks remained stable already from DIV 7 which is likely due to the unidirectional 2 node architecture. Due to missing reciprocal connections in this architecture, we expect synapse formation to have a lower impact on the overall spontaneous activity. The mean firing rate of around 14 Hz is comparable to previous publications using the same channel dimensions and microelectrode arrays (Ihle et al. (2022)) which is higher compared to random cultures because of the spatial confinement of axons (Pasquale et al. (2017)). The absolute value depends on the culture medium, neuron type, channel dimensions, the number of axons and the signal to noise ratio of the recording system. It is thus hard to compare between different studies (Chiappalone et al. (2006); Ihle et al. (2022)).

### Network integrity over varying channel lengths

Previous studies have explored axonal guidance and neuronal network formation, primarily focusing on short-range interactions (0.2-1 mm) (Isomura et al. (2015); Park et al. (2021); Winter-Hjelm et al. (2023); Mateus et al. (2021)). Retinal axons *in vivo* reach a total length of 2 cm, as apparent from the optic nerve in rats (Cavallotti et al. (2003)). Failures to cover the same distance *in vitro* are likely attributed to the experimental setup or to network viability. Axon viability in the long (6 and 8 mm) microchannels declined over time, as indicated by the significant decrease in the number of active electrodes during one month in culture. Since a decline was not observed in shorter channels, the reduced viability was most likely caused by channel length-related constraints. Millet *et al*. (Millet et al. (2007)) found similar trends when exploring the impact of channel length on cell viability in closed PDMS microstructures: Neurons in 3 mm long channels showed similar growth to unconstrained control cultures, whereas no viable cells were found in 6 mm long channels. Adding culture medium in a laminar flow to the closed PDMS channels enabled neurons to survive in up to 10 mm long (Millet et al. (2007)) channels, suggesting an impact of medium diffusion on cell viability that might also impact axon health (Walker et al. (2004); Morin et al. (2006)).

### Directionality of signal transmission

We achieved high directionality of axonal growth by combining directional edge guidance with a delayed target seeding strategy. Both methods have been used individually to achieve directionality rates of more than 90 % (Forró et al. (2018); Ming et al. (2021); Girardin et al. (2022); Pan et al. (2015)). For channels up to 2 mm, we achieved similar high values of forward propagating spikes (retina to thalamus) but observed a drop in directionality in longer channels. In longer channels, axons require more time to reach the target well compared to shorter channels, which might reduce the efficiency of the delayed seeding strategy (Pan et al. (2015)). To improve directionality in longer channels, the timepoint of target seeding could be increased with increasing channel length.

### Spike propagation speed

During neuronal maturation, the spike propagation speeds were shown to increase over time both *in vivo* (Foster et al. (1982)) and *in vitro* (Hong et al. (2016)). We observed a similar trend of increased spike propagation speed between 1 and 4 weeks *in vitro* in the shorter channels up to 4 mm but not in the 6 mm long channel. Since spike propagation speed depends on the axon diameter (Waxman and Swadlow (1977)), the most likely scenario is that axons in longer channels do not increase the average axon diameter in between the recording electrodes during 4 weeks in culture.

### Stimulation frequency and sustained neuronal response

The combination of electrical stimulation and calcium imaging enabled us to assess retinothalamic connectivity, and to distinguish presynaptic from postsynaptic activity (Moutaux et al. (2018)). The observation that most axons within the channel were from mRuby expressing RGCs, together with the high percentage of forward propagating spikes supports successful synaptic transmission. Moreover, previous publications investigating the functional strength between cortical neuronal populations suggested that about 100 axons are sufficient to induce postsynaptic bursts in the target well (Pan et al. (2015)) which is slightly more compared to the estimation of about 70 RGC axons within the presented system. However, our experiments could not exclude the possibility that a fraction of stimulation induced thalamic activity is the result of direct thalamic stimulation through back-growing axons.

Our results showed that a sustained target response could be elicited for stimulation frequencies between 4 and 31 Hz independent of the channel length. Stimulating at frequencies higher than 31 Hz resulted in a transient thalamic calcium response. Hadjinicolaou *et al*. showed that, in acute retinal recordings, the different retinal ganglion cell types showed varying maximum response frequencies with all RGCs having their cutoff between 10 and 70 Hz. Beyond this cutoff, the spiking response declined (Hadjinicolaou et al. (2016)). Similar studies using a directional two node network of cortical neurons showed that a 50-fold increase of the presynaptic stimulation frequency activated 4x more target neurons in dissociated cultures (Moutaux et al. (2018)). Thus, the maximum calcium response frequency in our culture does not only depend on the type and composition of retinal ganglion cell axons, but also on the number and type of thalamic target neurons. Consequently, it will be ultimately limited by the distribution and refractory period of the voltage sensitive sodium channels in each neuron (Wang et al. (2016)).

### Limitations

In this work, we have used primary rat neurons to establish the retinothalamic network model. The obtained results may not be translatable to other neuronal cell types such as human induced pluripotent stem cell (iPSC) derived neurons. *In vivo* human neurons cover greater distances and might show a higher growth potential compared to rat neurons. We cannot distinguish whether the limitation of 6 mm long channels is due to an intrinsic cell-growth limitation or to the decreased diffusion of nutrients in longer PDMS channels. Using bigger channels would increase nutrient diffusion but decrease the signal to noise ratio of recorded action potentials (Wang et al. (2012); Goshi et al. (2022)).

In this study, we used spheroids, which showed highly correlated neuronal activity within a single spheroid, thereby limiting the analysis on how individual neurons contribute to the compound calcium response at different stimulation frequencies.

## CONCLUSION AND OUTLOOK

In conclusion, our study shows the potential of *in vitro* systems to study retinogeniculate signal transmission. We show the feasibility of modeling unidirectional retinothalamic networks with up to 6 mm long axons and show the impact of axon channel length on network integrity and signal transmission fidelity. In future experiments, primary animal neurons may be replaced with human iPSC derived neuronal spheroids to study the effect of genetic or functional modifications on retinothalamic signal transmission and axon growth in healthy as well as diseased neurons (Fligor et al. (2021); Fernando et al. (2022)).

## AUTHOR CONTRIBUTIONS

GA, SI, JV and TR designed the experiments. GA conducted the experiments. GA, SJI and BC developed the python scripts and analysed the data. GA designed the PDMS microstructures. SG established and helped with spheroid formation. GA and TR wrote the first draft of the manuscript. BM and JD supported data analysis, interpretation and recording. All co-authors contributed to and approved the manuscript.

## FUNDING

The research was financed by ETH Zurich, the Swiss National Science Foundation (Project Nr: 165651), the Swiss Data Science Center, a FreeNovation grant, the OPO Foundation and the Human Frontiers Science Program Organization, HFSPO.

## Supplementary Material

### SUPPLEMENTARY FIGURES

**Figure S1.**
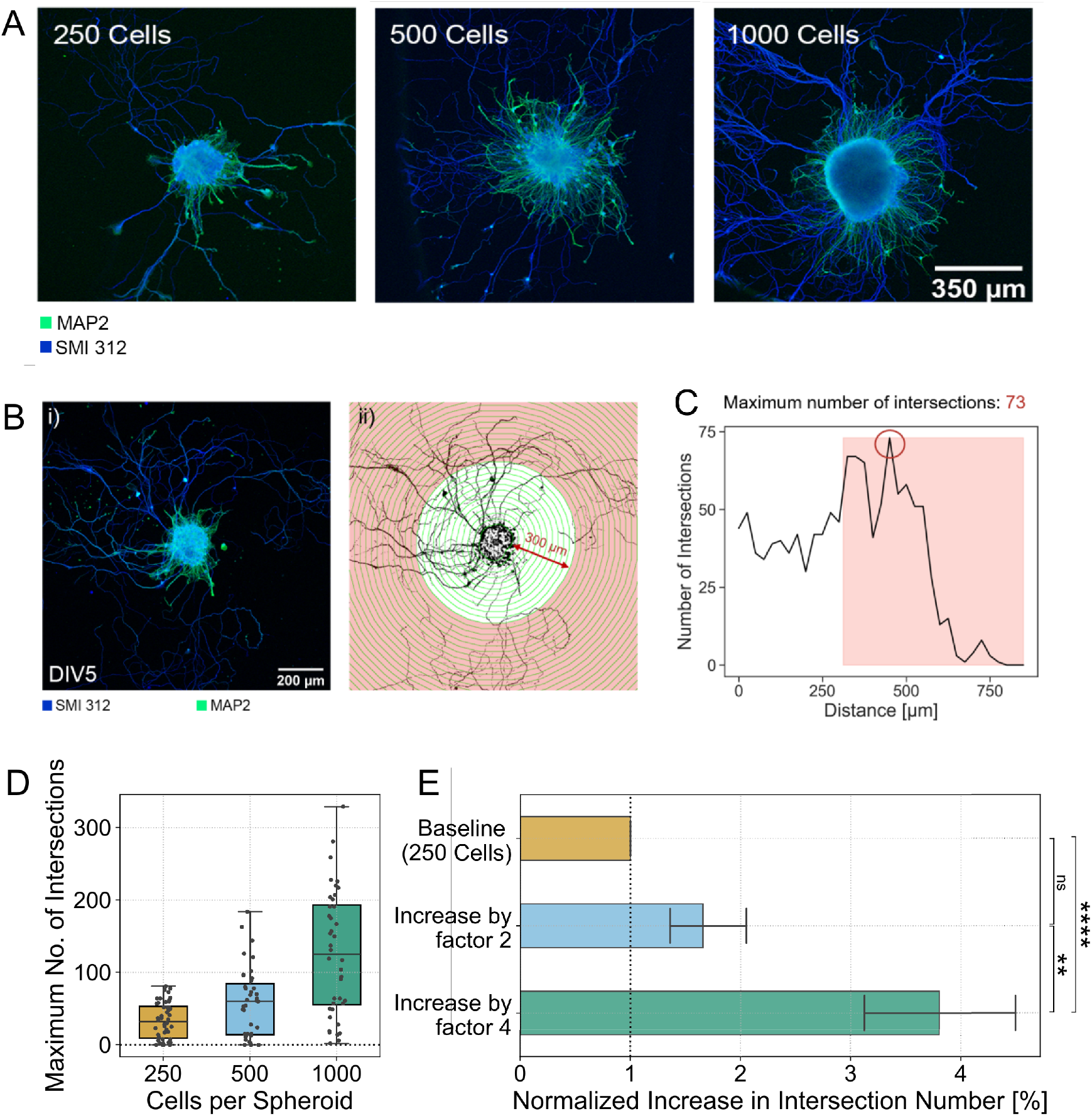
Correlation between retinal spheroid size and axon number. (**A**) Representative fluorescence images of retinal spheroids with different cell sizes: 250 cells/spheroid, 500 cells/spheroid, and 1000 cells/spheroid. Immunostaining was performed using antibodies against MAP2 (green, labels dendrites) and SMI 312 (red, labels axons). (**B**) Illustration of the performed Scholl analysis counting the number of intersections at a radius of 300 μm. This radius should exclude most dendrites. (**Ci**) Quantification of the maximum number of intersections at 300 μm. The number of intersections does not equal number of axons but gives an estimate of the minimum number of axons. (**Cii**) Relative increase in the number of intersections relative to 250 cell spheroids.

**Figure S2.**
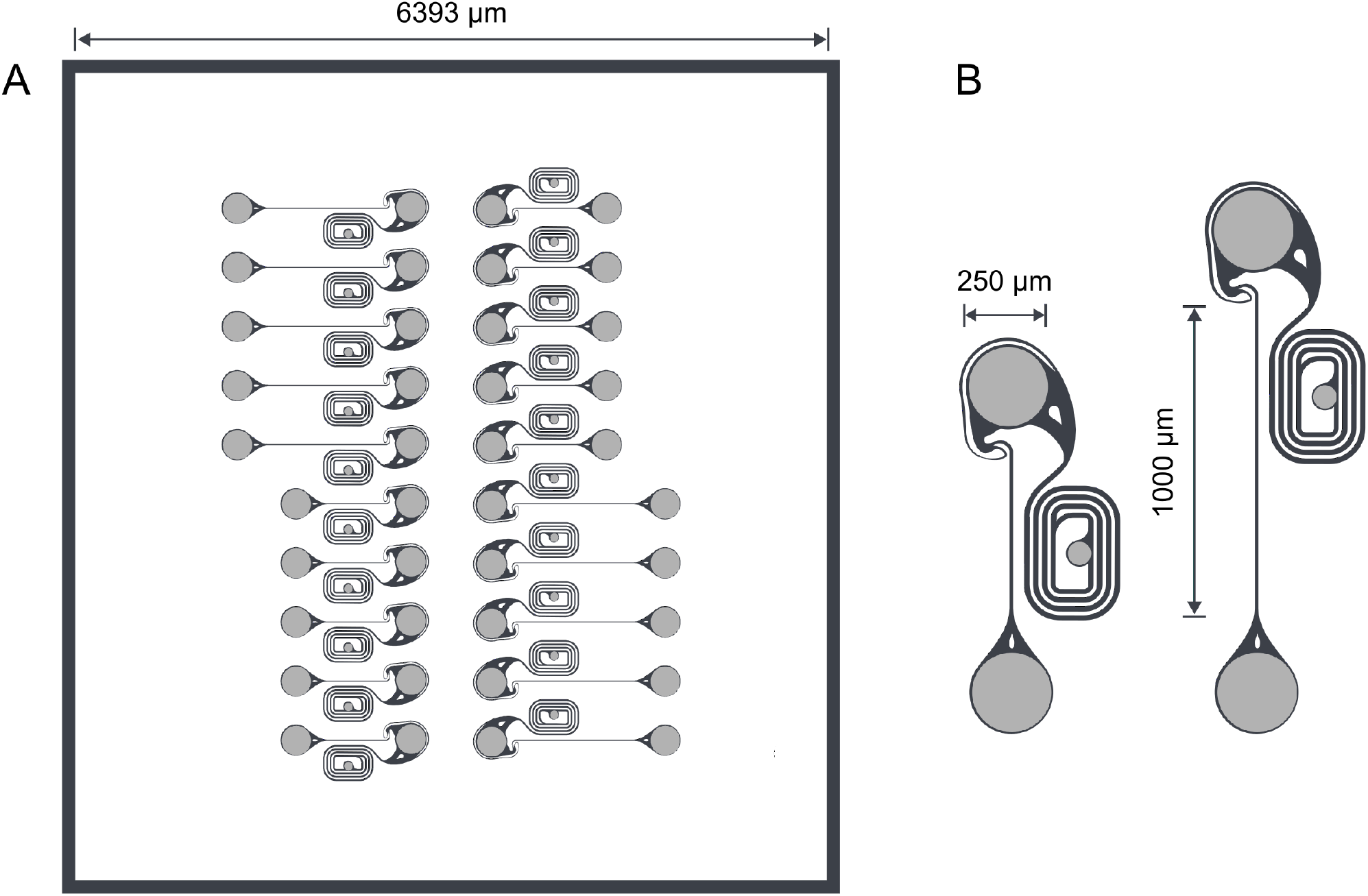
Microstructure designs with 1 and 0.5 mm channel length. (**A**) One microstructure fits 10 networks of the same channel length. (**B**) Only the channel length was varied between different microstructure architectures.

**Figure S3.**
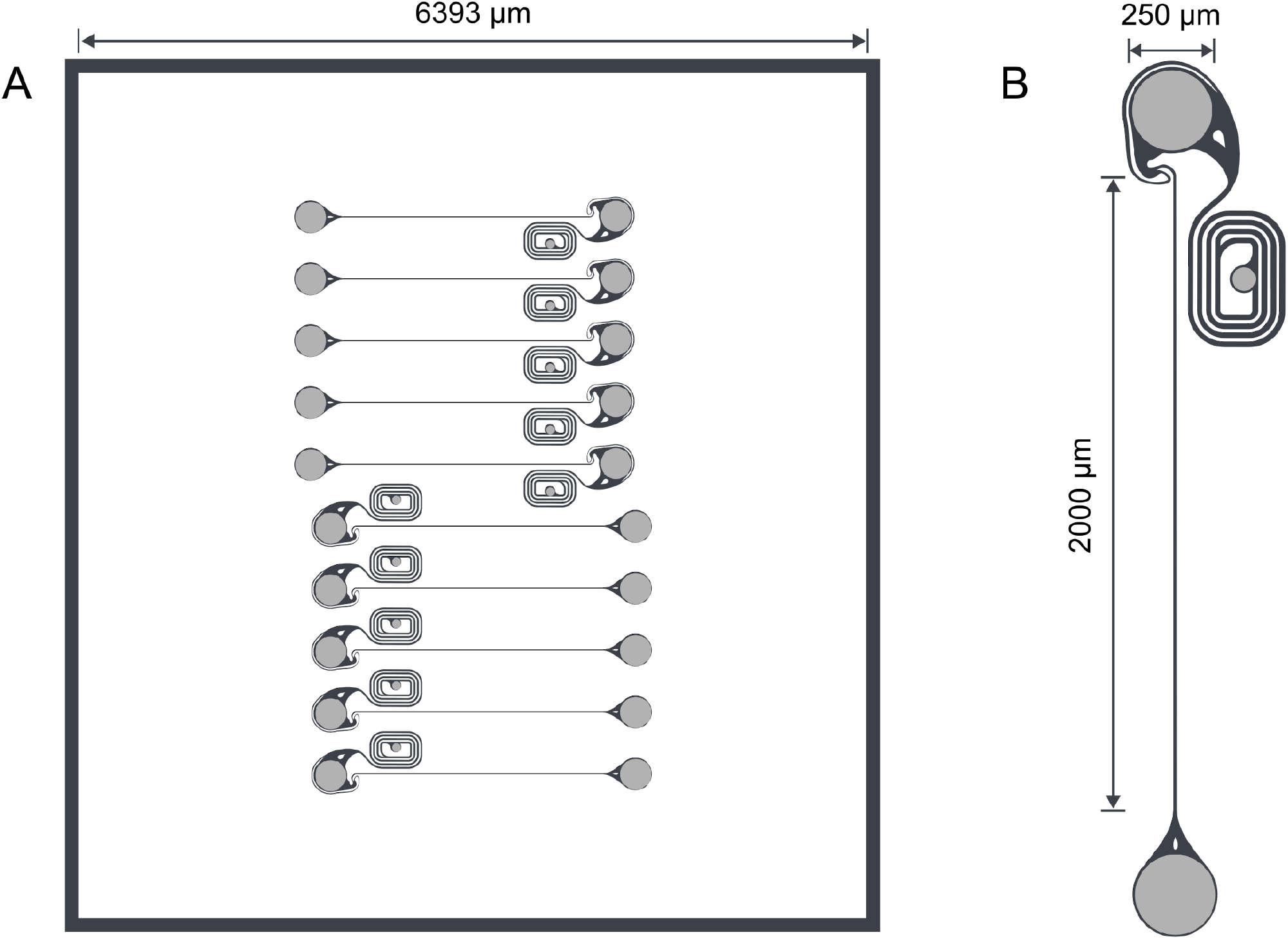
Microstructure designs with 2 mm channel length. (**A**) One microstructure fits 10 networks of the same channel length. (**B**) Only the channel length was varied between different microstructure architectures.

**Figure S4.**
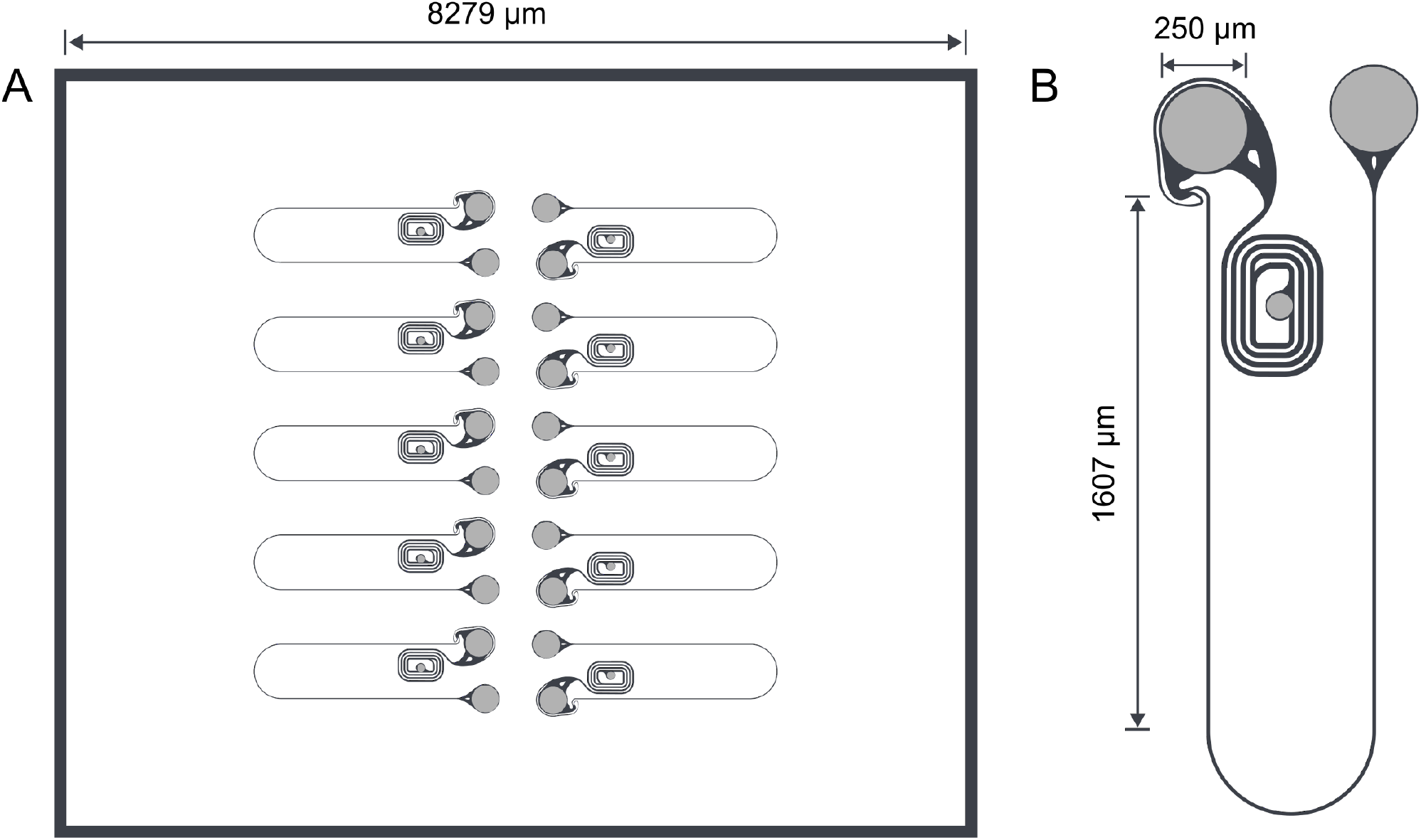
Microstructure designs with 4 mm channel length. (**A**) One microstructure fits 10 networks of the same channel length. (**B**) Only the channel length was varied between different microstructure architectures.

**Figure S5.**
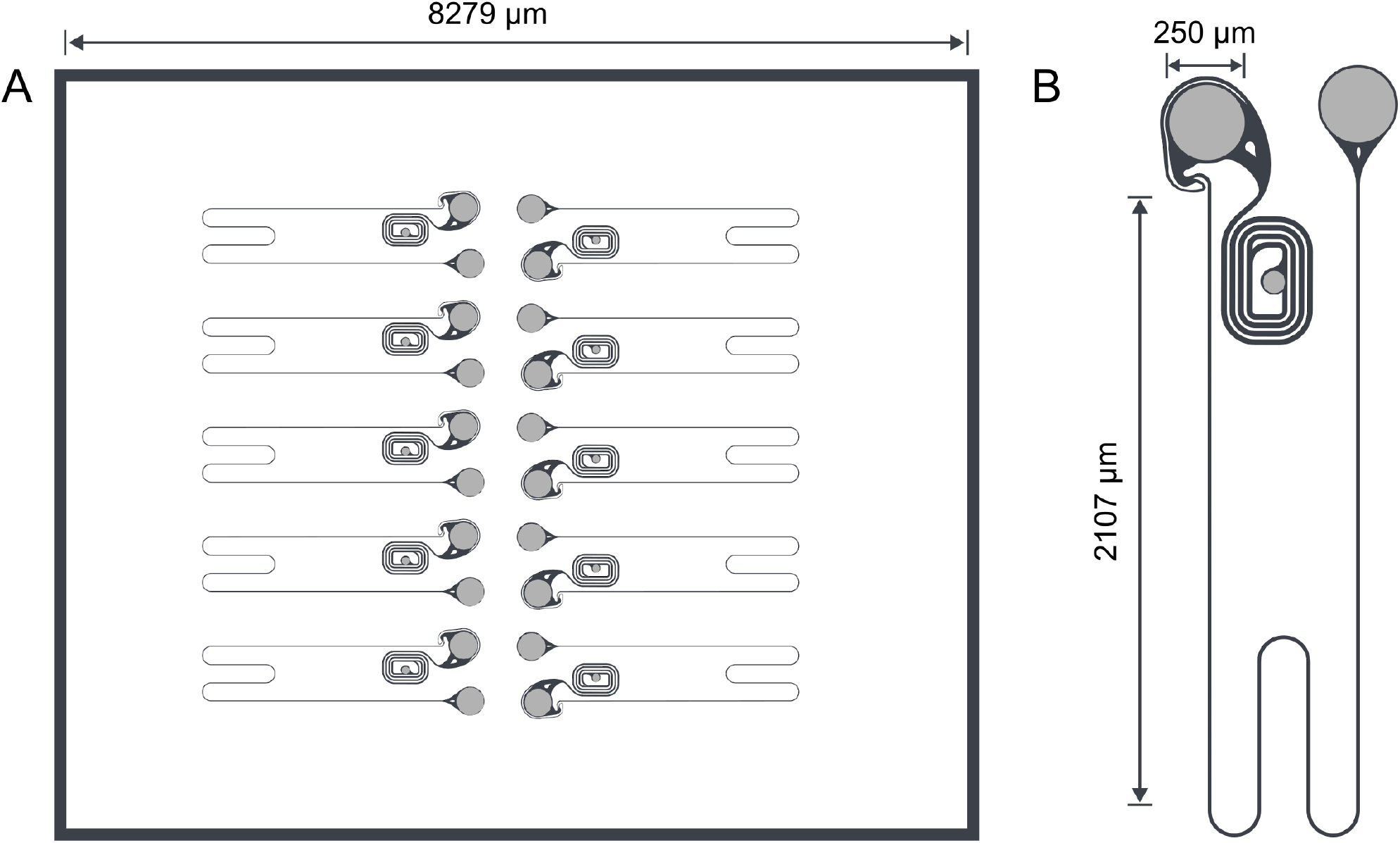
Microstructure designs with 6 mm channel length. (**A**) One microstructure fits 10 networks of the same channel length. (**B**) Only the channel length was varied between different microstructure architectures.

**Figure S6.**
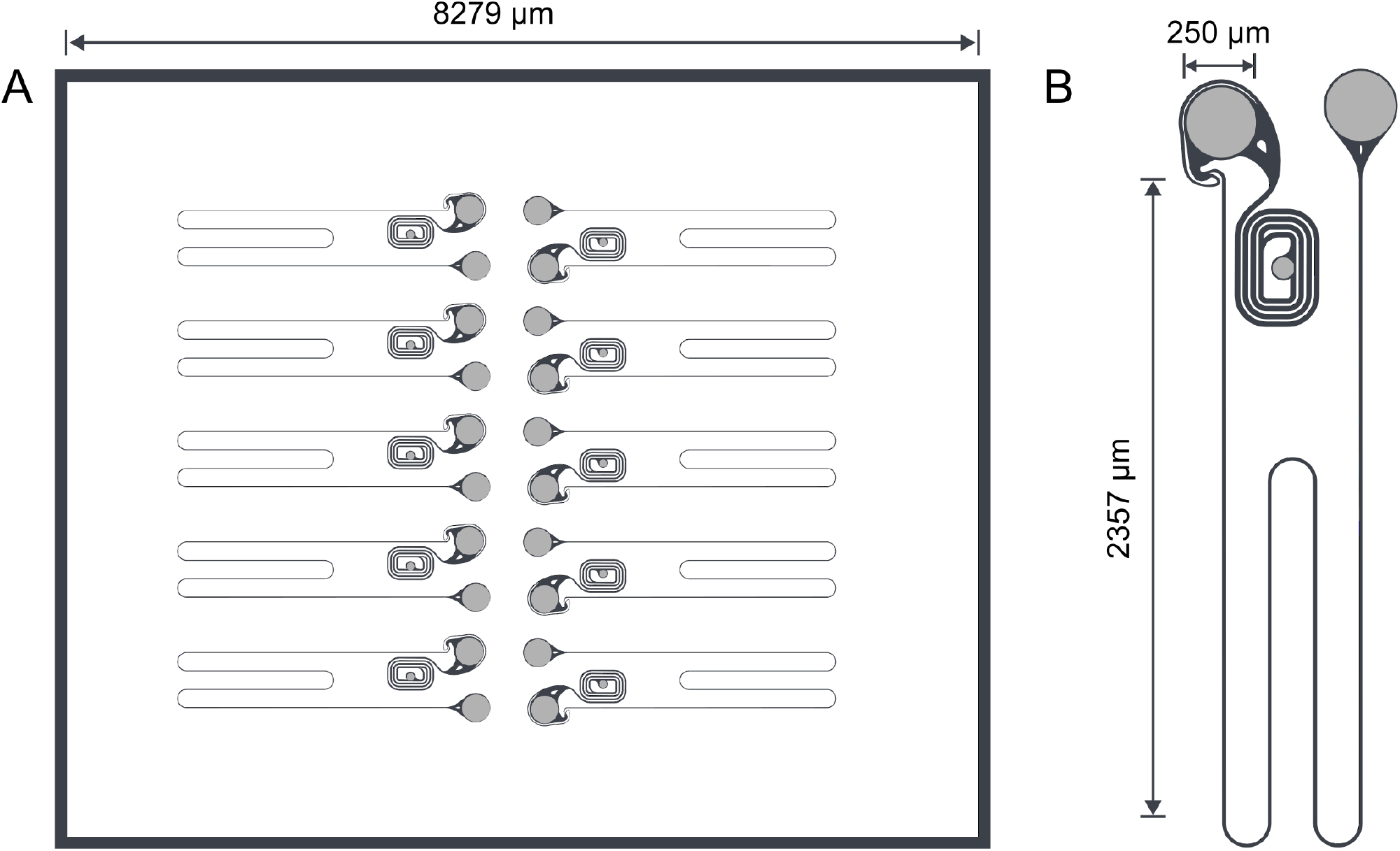
Microstructure designs with 8 mm channel length. (**A**) One microstructure fits 10 networks of the same channel length. (**B**) Only the channel length was varied between different microstructure architectures.

**Figure S7.**
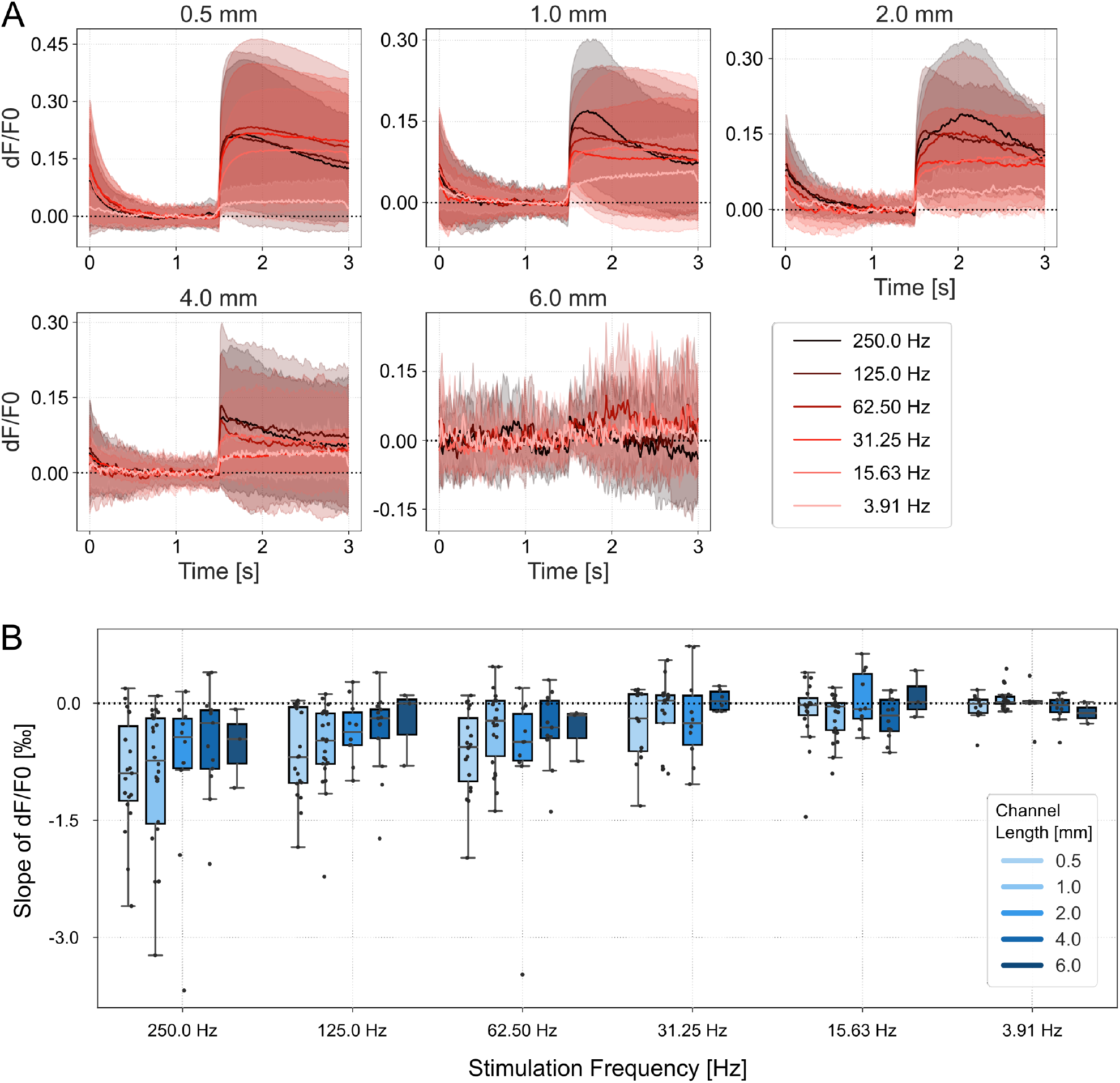
Thalamic response to continuous stimulation at varying frequencies. (**A**) Average calcium traces of thalamic spheroids in response to axonal stimulation at different frequencies over increasing channel lengths. Stimulus onset at 15 s. The areas represent the estimated mean with a 95 % confidence interval. Error; SEM. (**(B)**) Calcium signal during continuous stimulation. Negative values indicate a negative slope and thus a decrease in fluorescence intensity over the stimulation period. A value close to 0 indicates a sustained response. Error bars, SEM.

### SUPPLEMENTARY TABLES

**Table S1:**
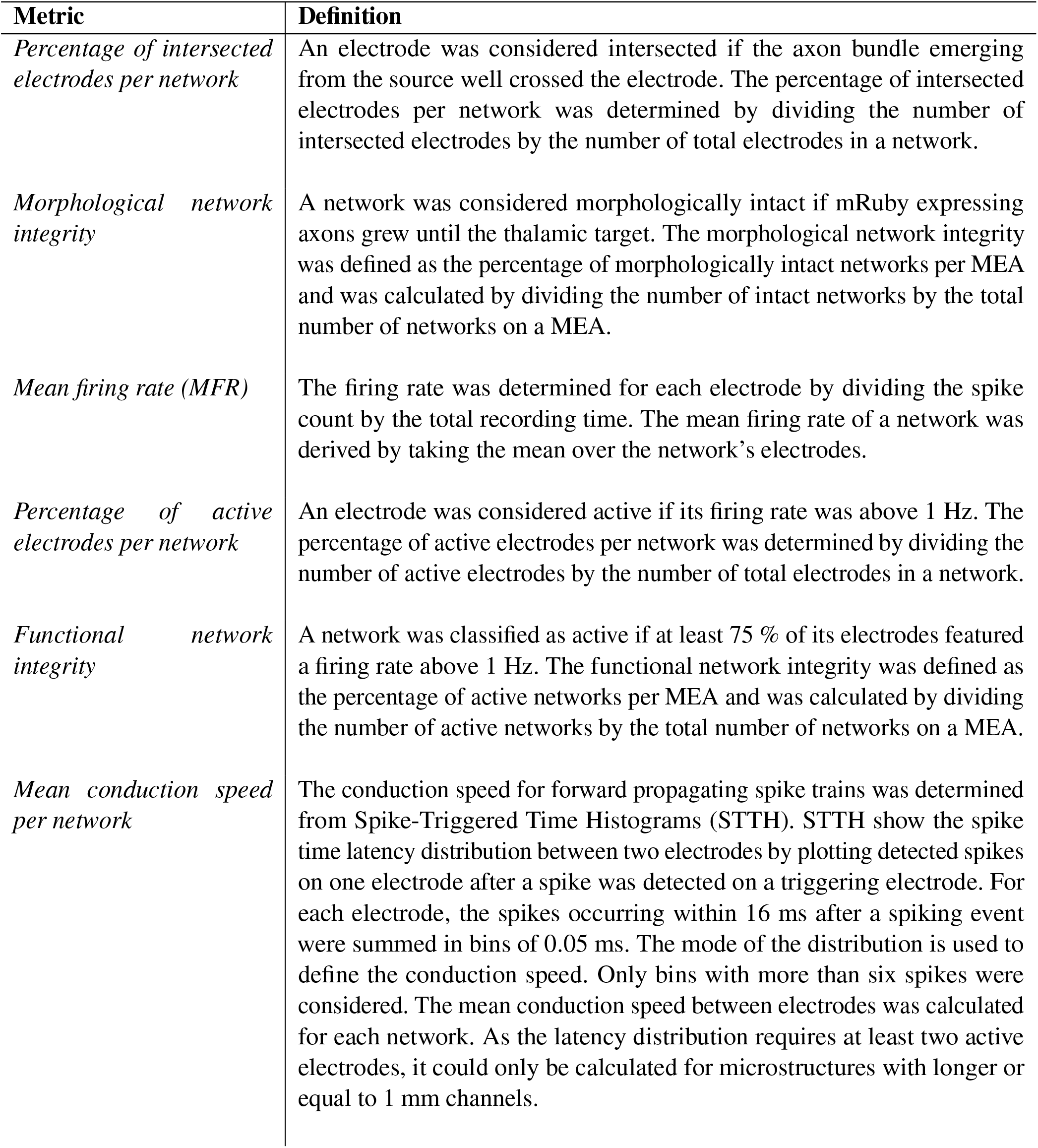
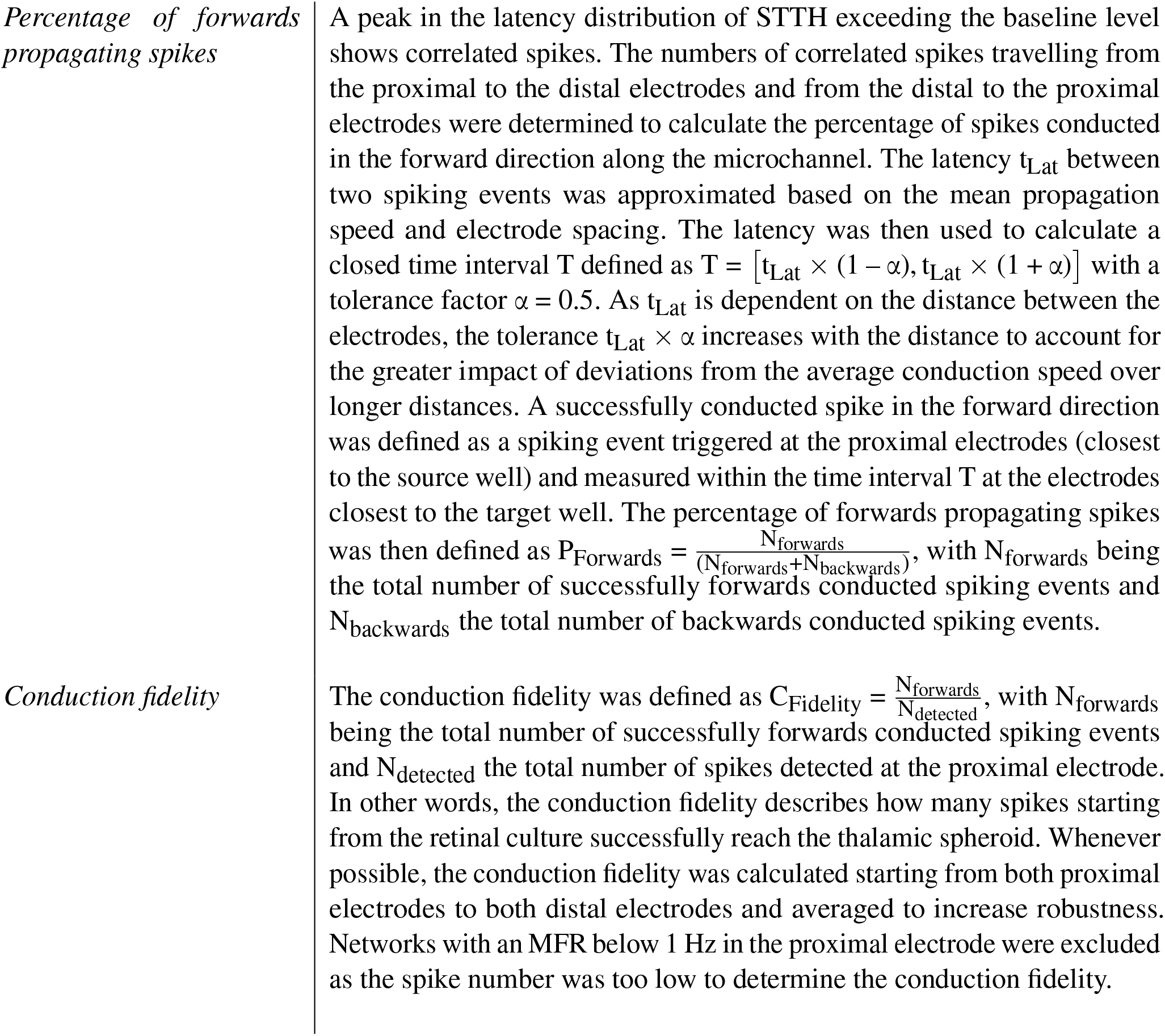
MEA metrics used to quantify the spike trains.

**Table S2.**
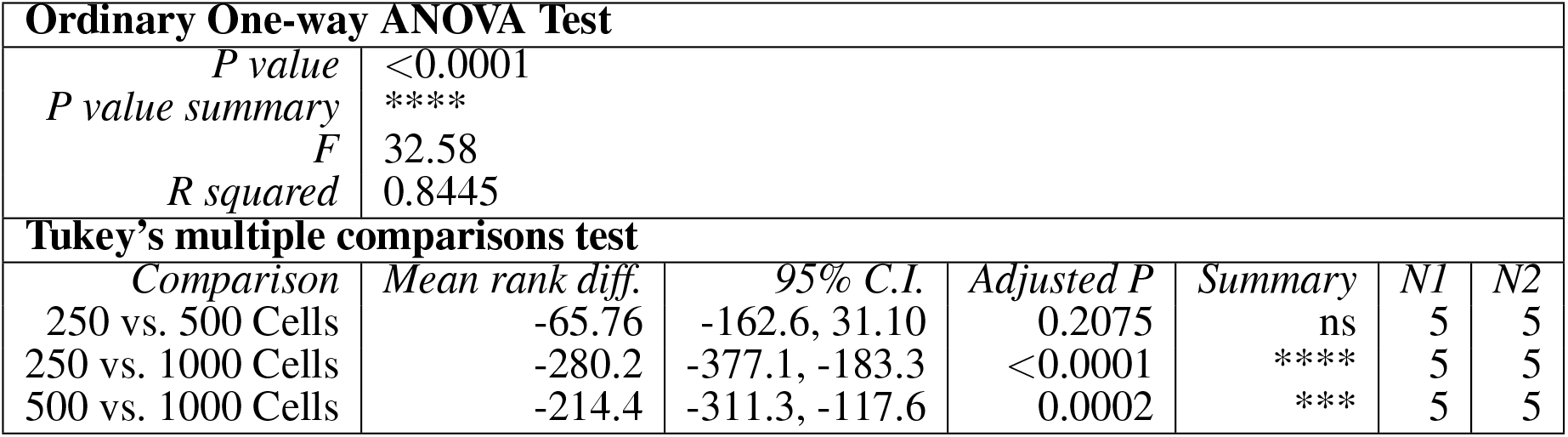
Ordinary one-way ANOVA test results for the axonal outgrowth data. Summary of the test and p-value statistics of the ANOVA test and Tukey’s multiple comparisons test for the relative increase in intersection number with spheroid size.

**Table S3.** Kruskal-Wallis test results for the percentage of active electrodes per network. Test and p-value statistics of the Kruskal-Wallis and post-hoc Dunn’s multiple comparisons test. The distributions of active electrodes per network were compared between the channel lengths.

| Kruskal-Wallis test |  |  |  |  |  |
| --- | --- | --- | --- | --- | --- |
| *P value* |  | <0.0001 |  |  |  |
| *Exact or approximate P value?* |  | Approximate |  |  |  |
| *P value summary* |  | **** |  |  |  |
| *Kruskal-Wallis statistic* |  | 115.9 |  |  |  |
| Dunn's multiple comparisons test |  |  |  |  |  |
| *Comparison* | *Mean rank diff.* | *Adjusted P Value* | *Summary* | *N1* | *N2* |
| 0.5 mm vs. 1 mm | -31.66 | >0.9999 | ns | 99 | 132 |
| 0.5 mm vs. 2 mm | -42.29 | >0.9999 | ns | 99 | 76 |
| 0.5 mm vs. 4 mm | 44.21 | >0.9999 | ns | 99 | 96 |
| 0.5 mm vs. 6 mm | 101.3 | 0.0003 | *** | 99 | 104 |
| 0.5 mm vs. 8 mm | 160 | <0.0001 | **** | 99 | 112 |
| 1 mm vs. 2 mm | -10.63 | >0.9999 | ns | 132 | 76 |
| 1 mm vs. 4 mm | 75.87 | 0.0133 | * | 132 | 96 |
| 1 mm vs. 6 mm | 132.9 | <0.0001 | **** | 132 | 104 |
| 1 mm vs. 8 mm | 191.7 | <0.0001 | **** | 132 | 112 |
| 2 mm vs. 4 mm | 86.5 | 0.0139 | * | 76 | 96 |
| 2 mm vs. 6 mm | 143.6 | <0.0001 | **** | 76 | 104 |
| 2 mm vs. 8 mm | 202.3 | <0.0001 | **** | 76 | 112 |
| 4 mm vs. 6 mm | 57.08 | 0.2666 | ns | 96 | 104 |
| 4 mm vs. 8 mm | 115.8 | <0.0001 | **** | 96 | 112 |
| 6 mm vs. 8 mm | 58.71 | 0.1692 | ns | 104 | 112 |

**Table S4.**
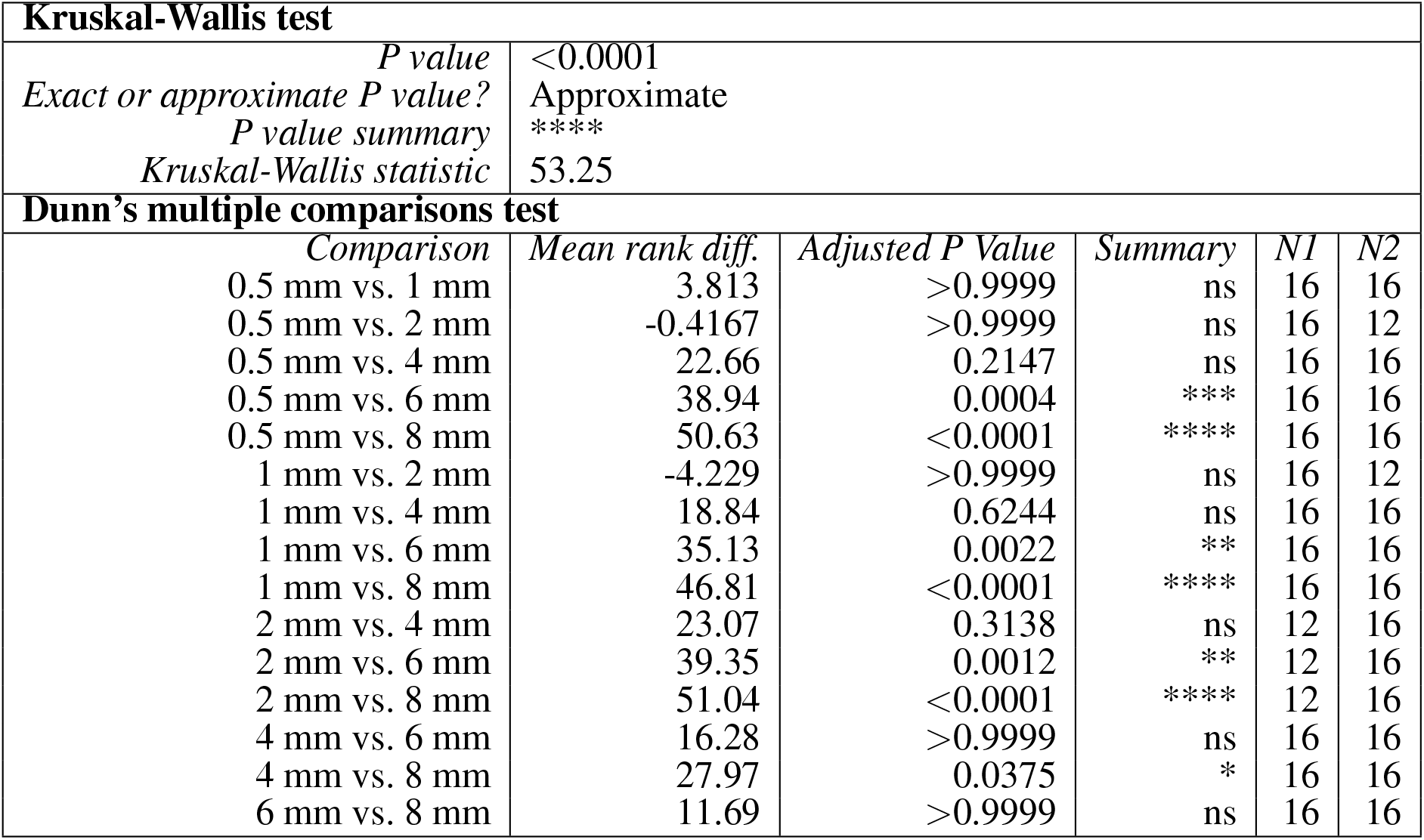
Kruskal-Wallis test results for the percentage of functionally intact networks. Summary of the test and p-value statistics of the Kruskal-Wallis and Dunn’s multiple comparisons test for the distributions of functional network integrity for each channel length.

**Table S5.**
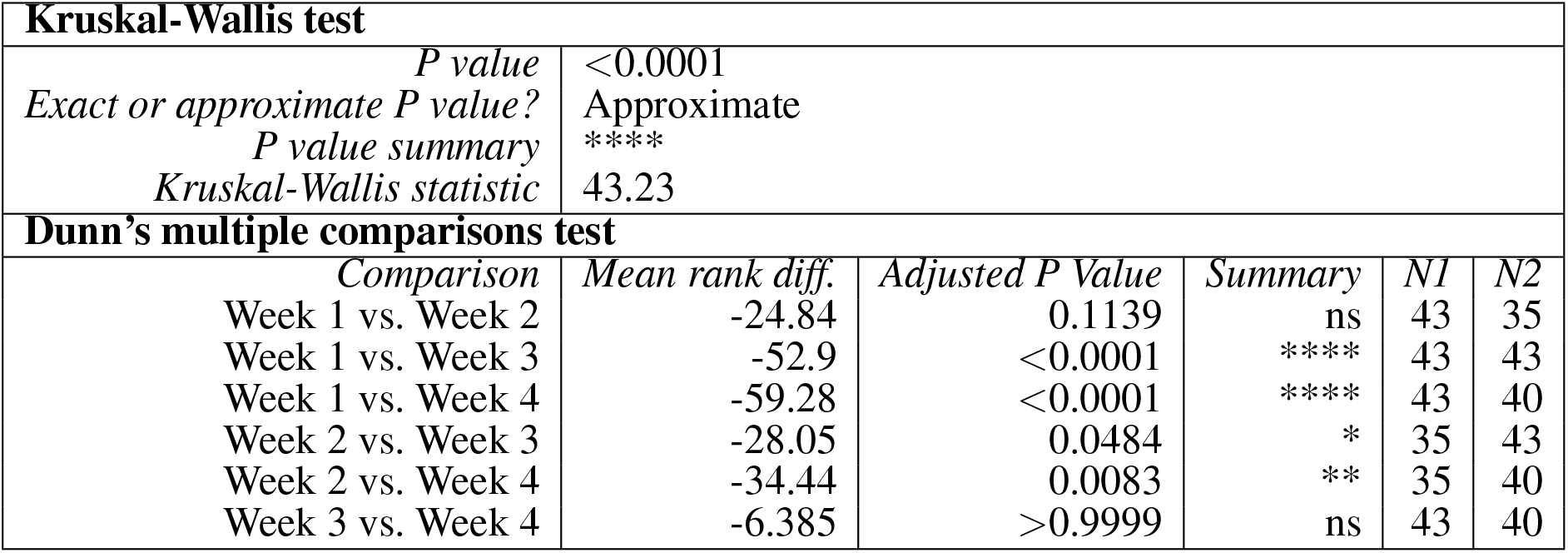
Kruskal-Wallis test results for the spike conduction speed over time. Test and p-value statistics of the Kruskal-Wallis and post-hoc Dunn’s multiple comparisons test. The conduction speed was compared between four weeks in culture.

**Table S6.** Friedman’s test results for the percentage of active electrodes per network over the weeks *in vitro*. Summary of the test and p-value statistics of the Friedman’s test and post-hoc Dunn’s multiple comparisons test. The distributions of active electrodes per network at different time points (week 1 - 4) were compared. The test was computed separately for each channel length.

| Friedman's Test for each channel lengths |  |  |  |  |  |
| --- | --- | --- | --- | --- | --- |
| Channel Length |  | Friedman statistic | P value | Summary | N |
| 0.5 mm |  | 1 | 0.8013 | ns | 25 |
| 1 mm |  | 1.941 | 0.5847 | ns | 33 |
| 2 mm |  | 0.2019 | 0.9773 | ns | 19 |
| 4 mm |  | 7.788 | 0.0506 | ns | 24 |
| 6 mm |  | 14.45 | 0.0024 | ** | 26 |
| 8 mm |  | 14.9 | 0.0019 | ** | 28 |
| Dunn's multiple comparisons test for the 6 mm long microchannels |  |  |  |  |  |
| Comparison | Rank sum diff. | Adjusted P Value | Summary | N1 | N2 |
| Week 1 vs. Week 2 | 5 | >0.9999 | ns | 26 | 26 |
| Week 1 vs. Week 3 | 16.5 | 0.458 | ns | 26 | 26 |
| Week 1 vs. Week 4 | 24.5 | 0.051 | ns | 26 | 26 |
| Week 2 vs. Week 3 | 11.5 | >0.9999 | ns | 26 | 26 |
| Week 2 vs. Week 4 | 19.5 | 0.2172 | ns | 26 | 26 |
| Week 3 vs. Week 4 | 8 | >0.9999 | ns | 26 | 26 |
| Dunn's multiple comparisons test for the 8 mm long microchannels |  |  |  |  |  |
| Comparison | Rank sum diff. | Adjusted P Value | Summary | N1 | N2 |
| Week 1 vs. Week 2 | 11.5 | >0.9999 | ns | 28 | 28 |
| Week 1 vs. Week 3 | 21 | 0.1784 | ns | 28 | 28 |
| Week 1 vs. Week 4 | 25.5 | 0.0498 | * | 28 | 28 |
| Week 2 vs. Week 3 | 9.5 | >0.9999 | ns | 28 | 28 |
| Week 2 vs. Week 4 | 14 | 0.8838 | ns | 28 | 28 |
| Week 3 vs. Week 4 | 4.5 | >0.9999 | ns | 28 | 28 |

**Table S7.**
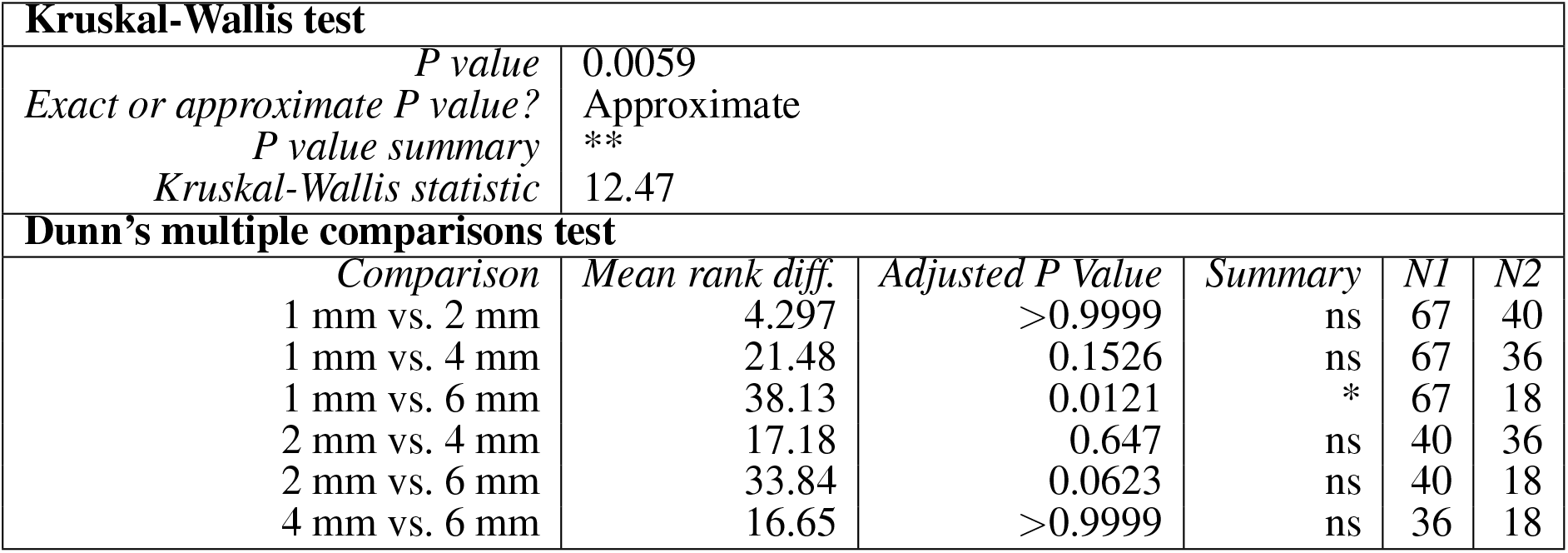
Kruskal-Wallis test results for the effect of channel length on spike conduction speed. Test and p-value statistics of the Kruskal-Wallis and post-hoc Dunn’s multiple comparisons test. The conduction speed was compared between neuronal networks in microstructures of different channel lengths.

**Table S8.**
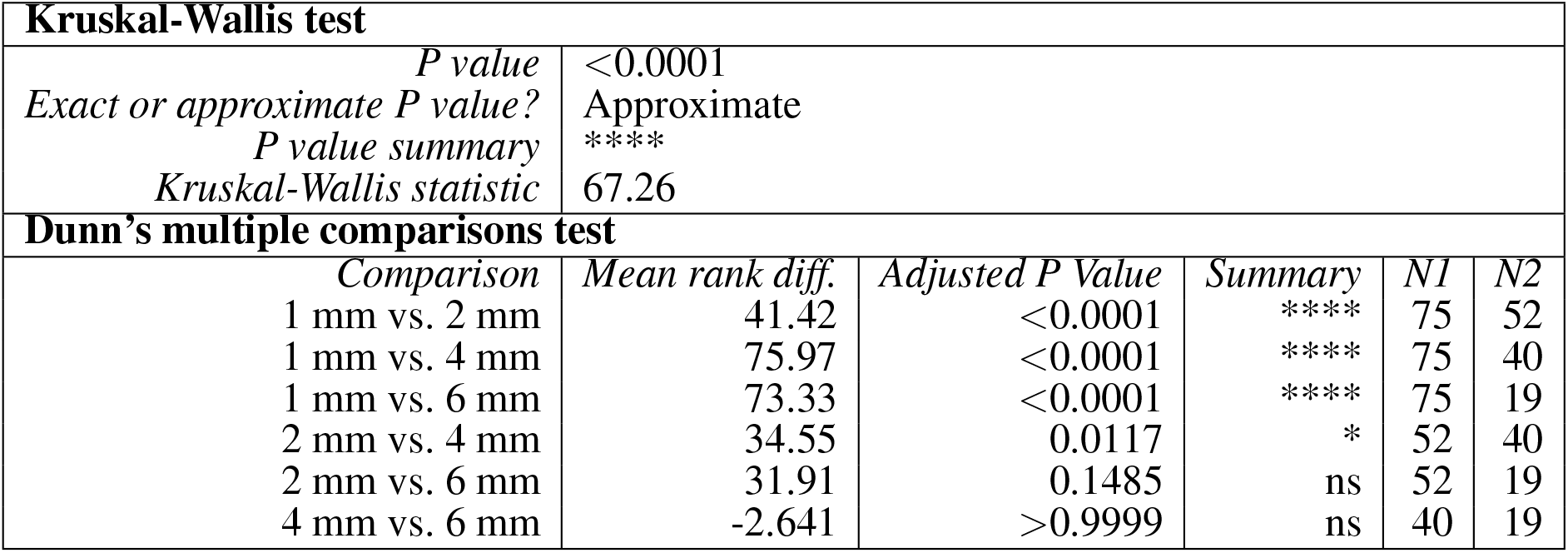
Kruskal-Wallis test results for the percentage of forwards propagating spikes in dependence on channel length. Test and p-value statistics of the Kruskal-Wallis and post-hoc Dunn’s multiple comparisons test. The percentage of forwards propagating spikes was compared between neuronal networks in microstructures of different channel length.

| Kruskal-Wallis test |  |  |  |  |  |  |
| --- | --- | --- | --- | --- | --- | --- |
| *P value* |  | <0.0001 |  |  |  |  |
| *Exact or approximate P value?* |  | Approximate |  |  |  |  |
| *P value summary* |  | **** |  |  |  |  |
| *Kruskal-Wallis statistic* |  | 67.26 |  |  |  |  |
| Dunn's multiple comparisons test |  |  |  |  |  |  |
| *Comparison* |  | *Mean rank diff.* | *Adjusted P Value* | *Summary* | *N1* | *N2* |
| 1 mm vs. 2 mm |  | 41.42 | <0.0001 | **** | 75 | 52 |
| 1 mm vs. 4 mm |  | 75.97 | <0.0001 | **** | 75 | 40 |
| 1 mm vs. 6 mm |  | 73.33 | <0.0001 | **** | 75 | 19 |
| 2 mm vs. 4 mm |  | 34.55 | 0.0117 | * | 52 | 40 |
| 2 mm vs. 6 mm |  | 31.91 | 0.1485 | ns | 52 | 19 |
| 4 mm vs. 6 mm |  | -2.641 | >0.9999 | ns | 40 | 19 |

**Table S9.**
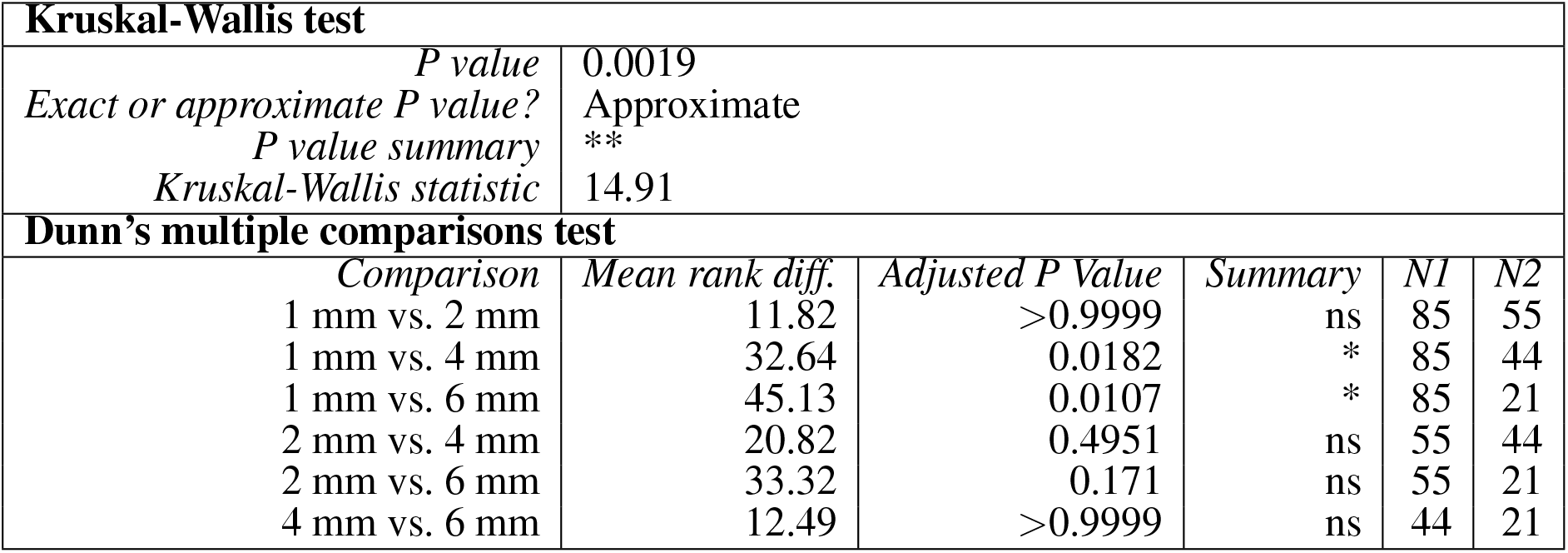
Kruskal-Wallis test results for the conduction fidelity in dependence on channel length. Test and p-value statistics of the Kruskal-Wallis and post-hoc Dunn’s multiple comparisons test. The conduction fidelity was compared between neuronal networks in microstructures of different channel length.

**Table S10.**
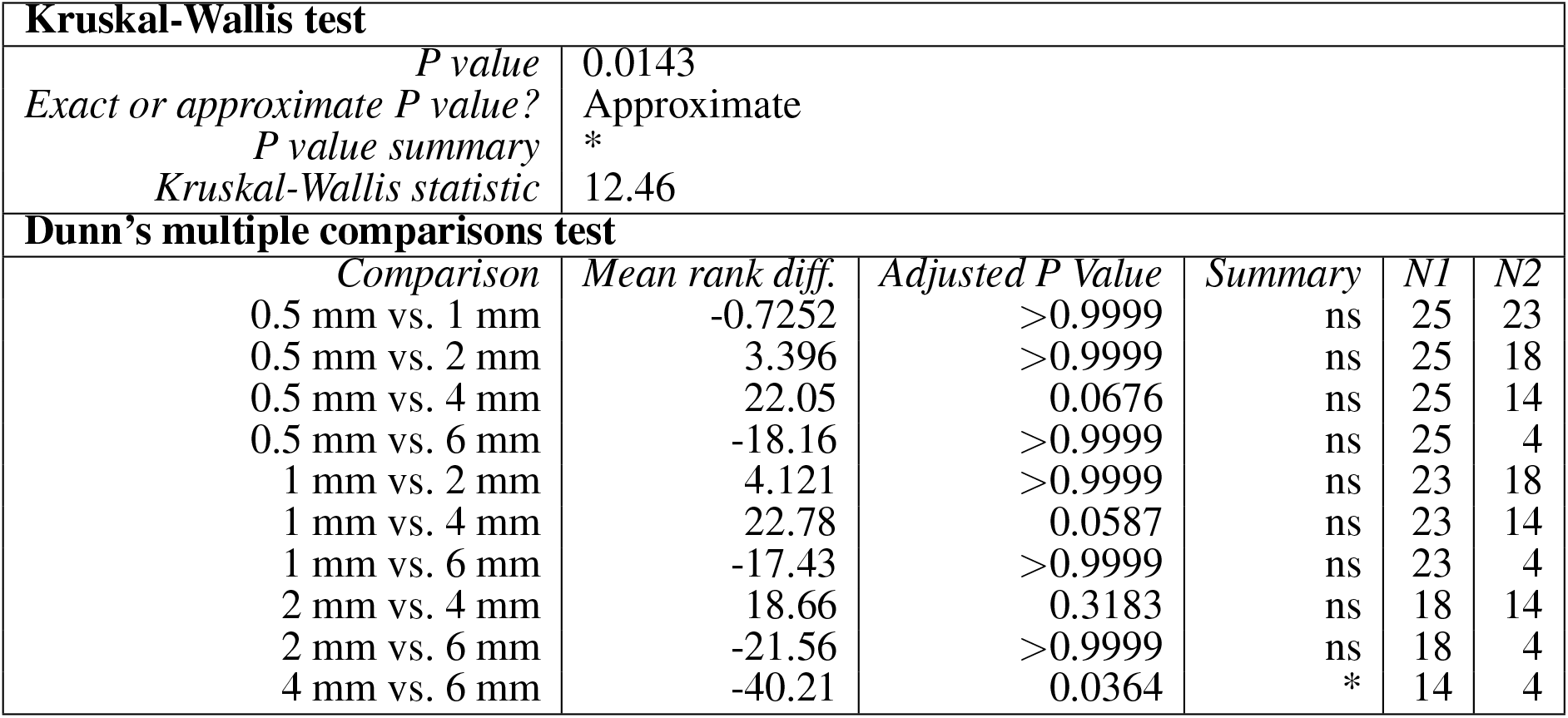
Kruskal-Wallis test results for the effect of the channel length on the peak evoked response. Test and p-value statistics of the Kruskal-Wallis and post-hoc Dunn’s multiple comparisons test. The PER was compared between neuronal networks in microstructures of different channel length.

## Notes

### Competing Interest Statement

The authors have declared no competing interest.

